# *Uhrf1* loss disrupts Ctcf-associated chromatin organization during early mouse embryogenesis

**DOI:** 10.64898/2026.07.15.738670

**Authors:** Agata Kurowski, Bohan Zhu, Yifei Sun, Paul Sargunas, Nimisha Jain, Ashley Chan, Lahouaria Hadri, Yi Shi, Martin J. Walsh, Sai Ma

**Affiliations:** Department of Pharmacological Sciences, Icahn School of Medicine at Mount Sinai, New York, NY 10029, USA; The Mount Sinai Center for RNA Biology and Medicine, Icahn School of Medicine at Mount Sinai, New York, NY 10029, USA; Department of Genetics and Genomic Sciences, Icahn School of Medicine at Mount Sinai, New York, NY 10029, USA; Institute of Genomic Health, Icahn School of Medicine at Mount Sinai, New York, NY 10029, USA; Department of Dermatology, Icahn School of Medicine at Mount Sinai, New York, NY 10029, USA

**Keywords:** UHRF1, DNA methylation, chromatin organization, embryonic development

## Abstract

UHRF1 is a chromatin-binding protein essential for maintaining DNA methylation and histone modification states, yet its integrated role *in vivo* remains incompletely understood. To define its function, we generated conditional *Uhrf1* knockout embryonic stem cells (ESCs) and embryos. *Uhrf1*⁻^/^⁻ ESCs exhibited near-complete loss of 5mC and 5hmC but maintained pluripotency, whereas Uhrf1-null embryos developed normally until E8.5 and then failed to develop further by E9.5, phenocopying *Dnmt1* loss. Single-cell multi-omic (ME-seq) profiling of E8.5 embryos revealed impaired lineage stabilization, widespread hypomethylation, and disrupted chromatin architecture. *Uhrf1* loss was associated with altered CTCF-associated chromatin signal, broad remodeling of chromatin contacts, altered *cis*-regulatory relationships, and reduced predicted BMP-related ligand–receptor communication, particularly within neural crest populations. These findings identify *Uhrf1* as a central regulator that tightly couples DNA methylation maintenance to 3D genome organization during gastrulation, thereby, directing early lineage specification and positioning *Uhrf1* as a pivotal mediator of epigenetic information transfer during early embryogenesis.

**Highlights:**

- *Uhrf1* knockout ESCs show global loss of 5mC/5hmC but maintain pluripotency.
- *Uhrf1*-null embryos develop normally until E8.5 but die by E9.5 with severe defects.
- Single-cell multi-omics revealed disrupted chromatin, transcription, and lineage stability upon knockout.
- Loss of Uhrf1 alters CTCF-associated chromatin signal, predicted BMP-related communication, and *cis*-regulatory relationships.

**Graphical abstract:** 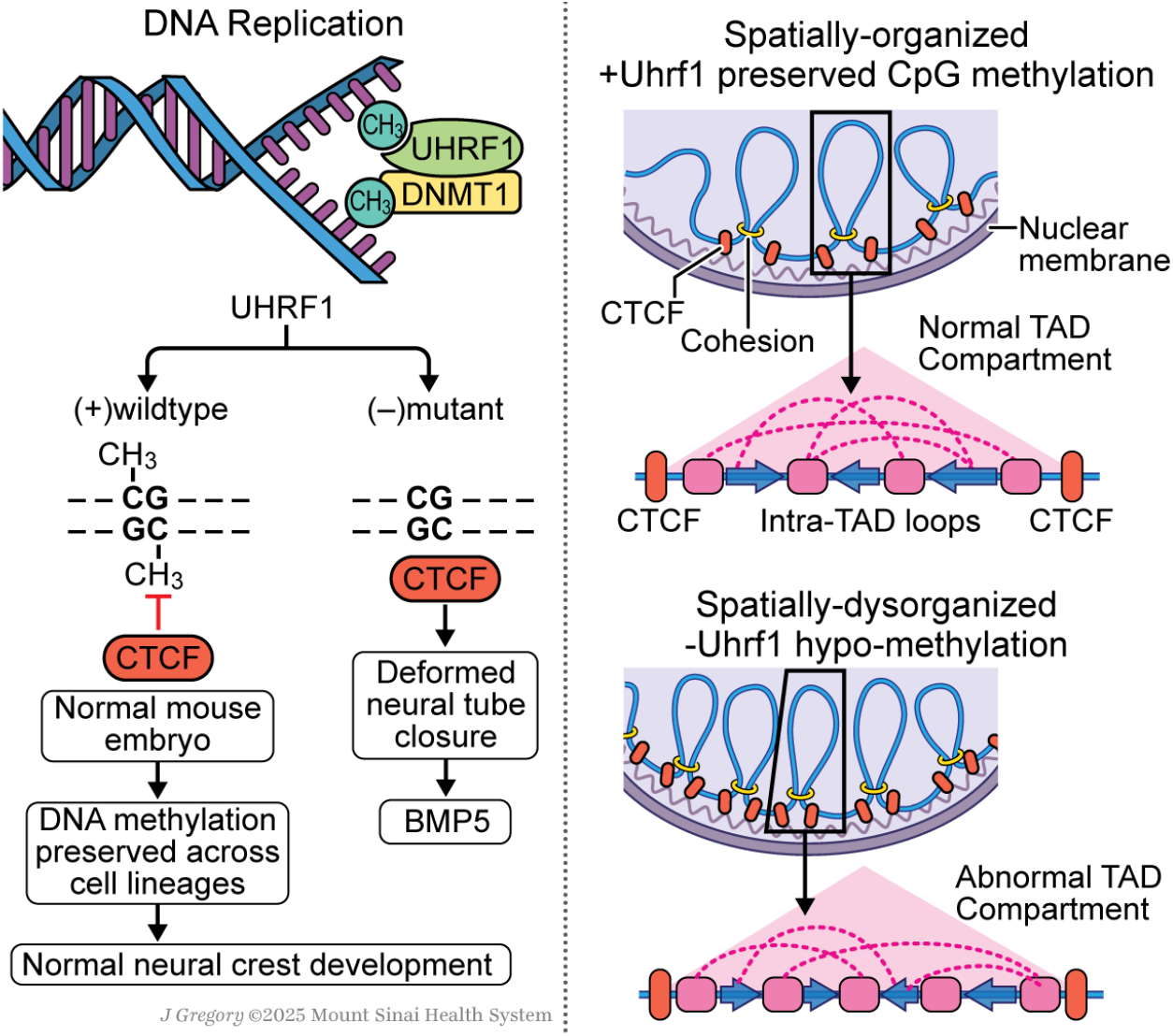

## Introduction

UHRF1 is an essential chromatin-binding protein that coordinates DNA and histone modification states to maintain epigenetic fidelity across cell divisions. Also known as ICBP90 in humans^1^ and NP95 in mice^2^, UHRF1 is composed of multiple reader and writer domains that recognize hemi-methylated DNA (via its SRA domain)^3^ and histone modifications (via its TTD and PHD domains)^4^, while its RING domain catalyzes H3 ubiquitination to recruit DNMT1 to ensure DNA methylation maintenance^5–7^. Through its interaction with G9a (KMT1C, EHMT2), the bivalent H3K4me1-K9me2/3 interaction of the tandem Tudor domain (TTD) and plant homeobox domain (PHD)^8^ influences chromatin binding of full-length UHRF1. This, in turn, localizes UHRF1 to enhancers and promoters of cell-type-specific genes at the flanks of cell-type-specific transcription factor binding sites, ensuring precise coordination of histone modifications and DNA methylation of those genes during S phase^9^. Additionally, UHRF1-dependent H3K23 ubiquitination couples maintenance DNA methylation and replication^6^.

Together, these mechanisms would be consistent with previous studies detailing the fundamental role of G9a, in concert with UHRF1, in maintaining DNA methylation, retrotransposon silencing, and chromatin topology during S-phase processes with UHRF1^10^. Through these multifaceted interactions, UHRF1 serves as a key factor linking DNA methylation, histone modification, ubiquitination, and chromatin organization into an integrated epigenetic network that ensures faithful inheritance yet can be co-opted in disease.

In cancer, UHRF1 is frequently overexpressed, particularly in highly proliferative tumors such as breast, lung, colorectal, and liver cancers, where it can disrupt DNA methylation, causing focal hypermethylation and thereby silencing tumor suppressor loci^11^. As noted previously^12,13^, the weakened capacity for CTCF to bind its cistrome when methylated cytosines are present, the loss of UHRF1’s epigenetic activity would interfere with CTCF occupancy by altering CpG methylation at boundary elements, thereby compromising topologically associating domain (TAD) structure and fidelity of enhancer architecture^14,15^. These findings provide a basis for UHRF1’s role in gene silencing and chromatin topology dysregulation, positioning it as both a biomarker and a mechanistic driver of tumorigenesis. Targeting UHRF1’s reader or ligase domains may offer therapeutic potential in restoring epigenetic homeostasis and chromatin architecture in cancer.

Although individual domain functions are well described, how UHRF1 functions as a regulatory factor that integrates these key fundamental biochemical and epigenetic roles *in vivo* remains poorly understood. Here, we examined the function of UHRF1 in murine embryonic stem cells (ESCs) *in vitro*. Despite a global loss of 5mC and 5hmC upon UHRF1 deletion, ESCs retained pluripotency and exhibited no overt phenotype, possibly reflecting their adaptive transcriptional plasticity^16^. In contrast, this study reveals that UHRF1-null embryos developed normally until E8.5 but terminated development by E9.5, phenocopying DNMT1 deficiency^17^. Multi-omic profiling (scRNA-seq, scATAC-seq, and sc-methyl-seq) of E8.5 embryos revealed widespread chromatin and transcriptional dysregulation, implicating UHRF1 as a key regulator of early lineage stabilization and chromatin integrity during gastrulation.

## Results

### UHRF1 knockout leads to global hypomethylation and alters imprinted genes *in vitro*

To investigate the role of UHRF1 in development, we generated a mouse model with exons 6-10 of *Uhrf1* floxed (**Extended Data Fig.1A**). We derived ESCs and utilized tamoxifen-inducible CRE-recombination to obtain UHRF1^-/-^ ESCs. *Uhrf1* levels were confirmed to be depleted by a Western blot (**Extended Data Fig.1B**). UHRF1^-/-^ cells cultured in 2i/LIF^18^ showed no visible phenotype (**Extended Data Fig.1C**) and no changes in cell-cycle distribution compared to *Uhrf1*^fl/fl^ ESCs (**Extended Data Fig.1D**). RT-qPCR showed no change in the expression levels of pluripotency factors *Oct4* and *Nanog* (**Extended Data Fig.1E**). The pluripotency of the ESCs was confirmed by injecting WT and KO ESCs into the flanks of immunocompromised mice, which generated 4/4 teratomas from wild-type ESCs and 4/5 from UHRF1^-/-^ ESCs, indicating no impairment in contributing to teratoma formation. All three germ layers were detected in teratomas from UHRF1^-/-^ ESCs, indicating that in this environment UHRF1 is dispensable for their formation (**Extended Data Fig. 1F**). RNA-seq analysis confirmed the depletion of *Uhrf1* expression and did not show significant changes in pluripotency marker gene expression and differentiation-related genes. Interestingly, it revealed bidirectional changes of multiple imprinted genes, including an up-regulation of *Xlr3c* (**Extended Data Fig.1G, Supplemental Table 1**). As UHRF1 is known to given its association with DNA methylation maintenance^19^, we next investigated the methylomic changes upon UHRF1 knockout. Indeed, we confirmed a drastic reduction of global 5-methylcytosine (5mC) of UHRF1^-/-^ ESCs compared to wild-type ESCs using dot-blot analysis (**Extended Data Fig.1H**). Further, we performed whole-genome bisulfite sequencing (WGBS) of wild-type and UHRF1^-/-^ ESCs and found that only a marginal amount of methylation remained in UHRF1^-/-^ ESCs (**Extended Data Fig.1I**). The loss of UHRF1 in ESCs leads to an almost complete depletion of 5mC. Surprisingly, neither the expression of pluripotency-related genes nor differentiation-related genes seems to be affected by this loss, and UHRF1^-/-^ ESCs show no gross phenotype. Fast proliferating ESCs, like cancer cells, provide a model that can rapidly adapt to environmental changes and possibly mask phenotypes that would occur *in vivo*. For this reason, we next investigated UHRF1 in mouse embryo development *in vivo*.

### *Uhrf1* knockout is lethal with vast defects arising at E9.5

The covalent, reversible, and heritable modification of chromatin is a regulatory mechanism that allows for rapid changes in gene expression in response to cellular cues. The process of embryonic development relies on rapid and precise changes in gene expression programs to allow cells to differentiate in a timely manner into various cell lineages. Whereas maternal NP95/UHRF1 seems essential for zygotic to maternal transition^20–22^, zygotic UHRF1^-/-^ knock-out embryos survive until gastrulation^19^. The effects of UHRF1 ablation and the phenotypes and lineages affected have not been studied in detail. Using immunofluorescence, we confirmed the absence of UHRF1 protein expression in the blastocyst (Embryonic time (E) 3.5) of our mouse model (**Fig.1A**) and loss of *Uhrf1* RNA expression by bulk RNA-seq at E9.5. RNA-seq analysis did not identify significant changes in pluripotency genes and differentiation-related genes. Consistent with prior reports that Uhrf1-null embryos fail during early post-implantation development^19^, UHRF1-null embryos exhibited developmental arrest and lethality beyond E8.5. We found that *Uhrf1*^-/-^ embryos show no visible phenotype until E8.5, but severe defects arise at E9.5 (**Fig.1B**), with *Uhrf1*^-/-^ embryos not recovered after E9.5 (**Supplemental Table 2**). Specifically, at E9.5, we detected an increased level of apoptosis marker Cleaved-Caspase 3 in the head area of the neuroepithelial lining of the hindbrain, the facial-acoustic neural crest complex, and the auditory vesicle (**Fig.1C, D, Extended Data Fig.1J**), whereas no increased cell death was detected at E8.5.

**Figure 1.**
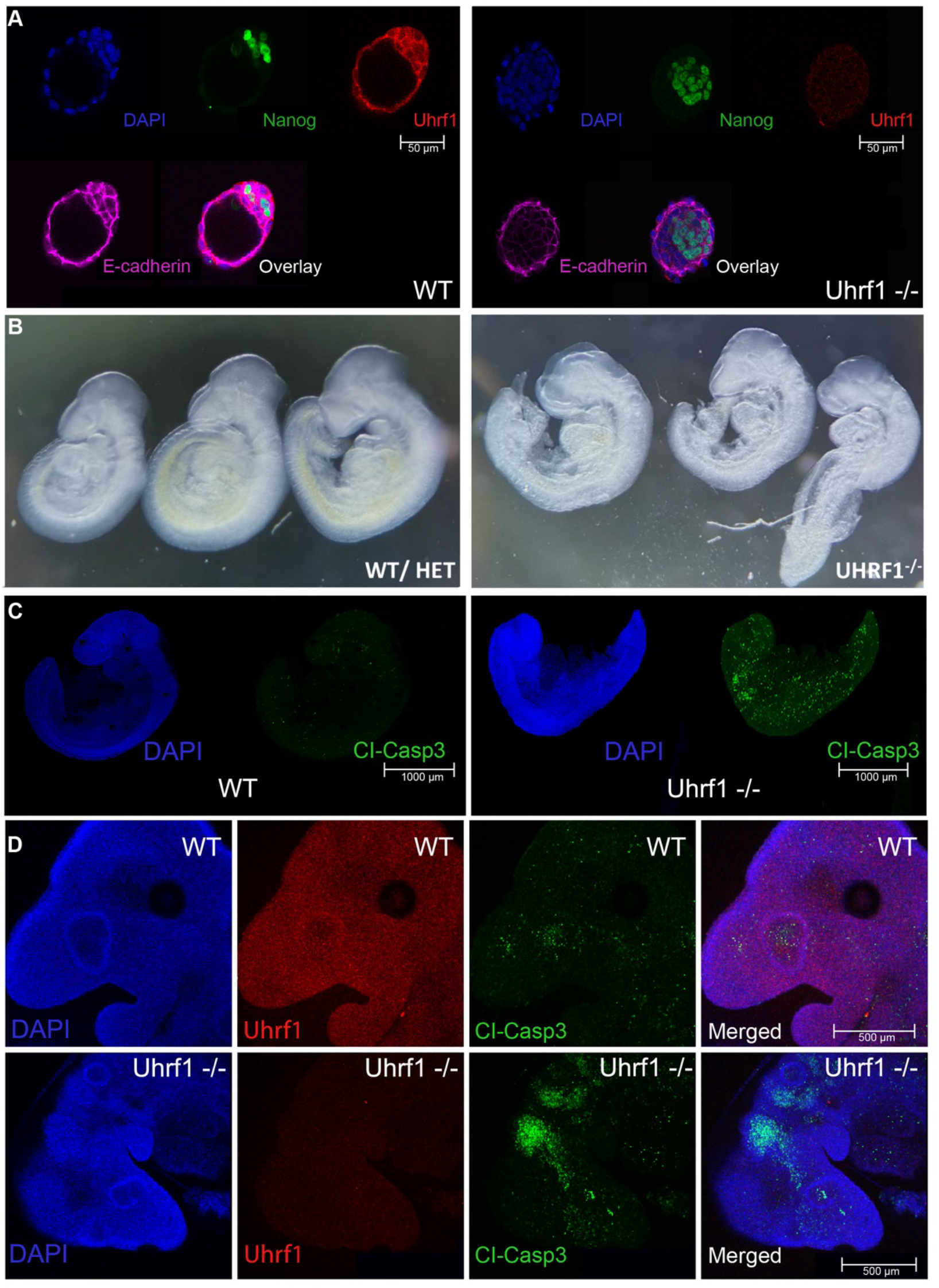
***Uhrf1* knockout is lethal with vast defects arising at E9.5 (A)** Absence of UHRF1 protein expression in E3.5 embryos. **(B)** Microscopic image of Uhrf1^-/-^ and wild-type E9.5 embryos. **(C,D)** E9.5 Uhrf1^-/-^ embryos show increased CI-Casp3 signal associated with cell death.

Similar to UHRF1^-/-^ embryos, DNMT1^-/-^ embryos die around gastrulation^23^. The lack of proper DNA methylation maintenance may be one cause of lethality. Given that UHRF1 has numerous additional functions on histone, DNA methylation, and gene regulation, we next examined UHRF1^-/-^ embryos at E8.5 by a single-cell multi-modal assay, ME-seq, which simultaneously captures transcriptome (scRNA-seq), open chromatin (scATAC-seq), and DNA methylation (scDNAme-seq), to investigate the cause of gene and chromatin dysregulation that leads to the decay of the embryo at E9.5.

### ME-seq captures diverse cell states in mouse embryos across three modalities

To simultaneously capture the DNA methylation status along with chromatin accessibility and gene expression changes, we leveraged ME-seq, which is built upon the SHARE-seq protocol^24^ (**Fig.2A**). Briefly, fixed cells isolated from four wild-type and four *Uhrf1* knockout E8.5 embryos were transposed with Tn5 to mark open chromatin. mRNA was reverse transcribed using a poly(T) primer containing a biotin tag; cells were distributed in a 96-well plate to hybridize well-specific barcoded oligonucleotides to chromatin and cDNA; hybridizations were repeated three times, expanding the barcoding space to ∼1×10^6^ (96^3^) which is sufficient to label every nucleus with a unique combination of barcodes; after reverse crosslinking, cDNA was separated from chromatin using streptavidin beads before downstream library preparation. The methylation status of DNA fragments was determined by converting methylated cytosine to dihydrouracil (DHU)^25^.

**Figure 2.**
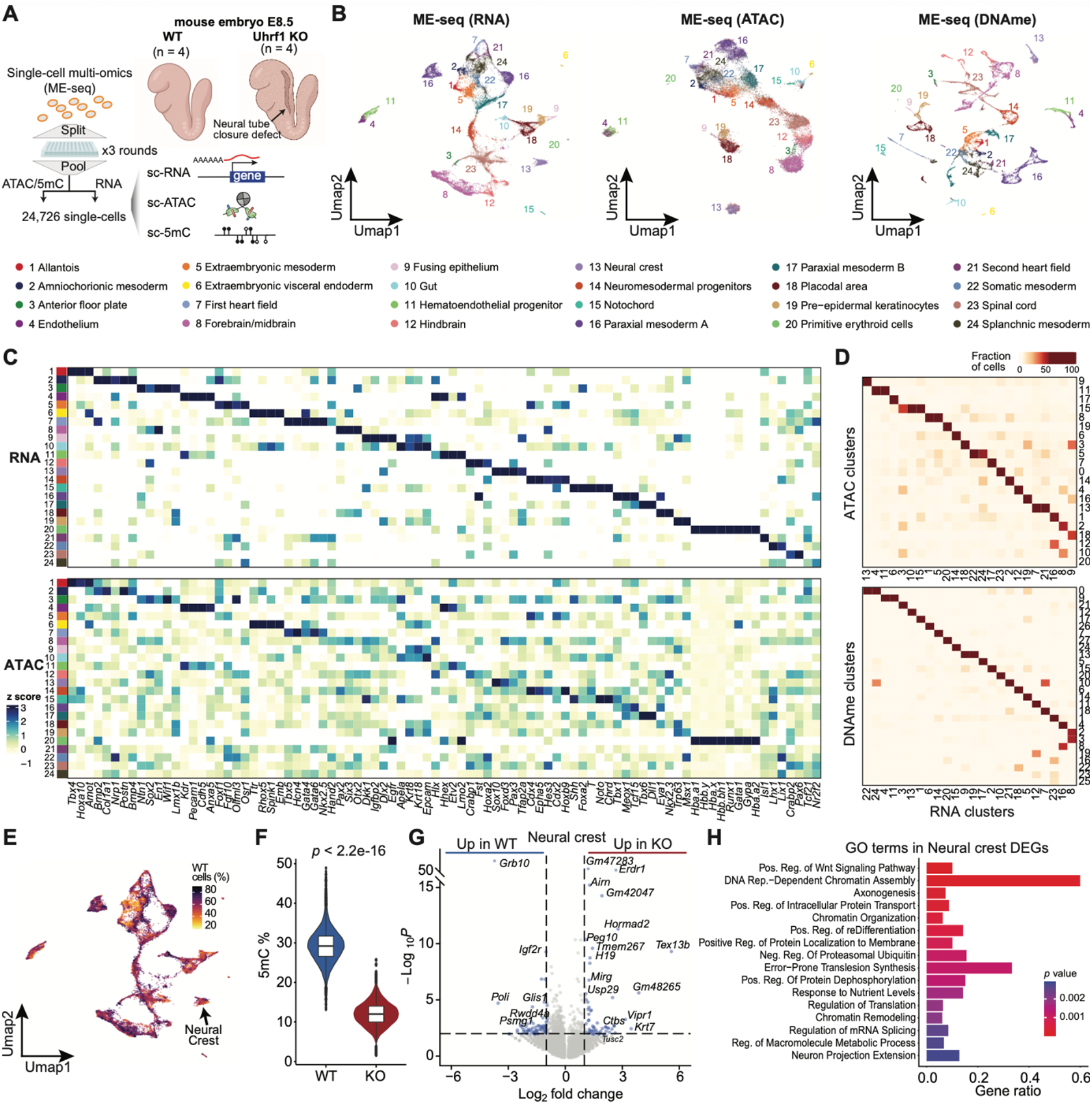
ME-seq captures diverse cell states in mouse embryos across three modalities. **(A)** Schematic of the experimental design. **(B)** ME-seq uniform manifold approximation and projection (UMAP) visualization of single cells derived from mouse embryos showing UMAP coordinates defined by RNA, ATAC, or DNA methylation. **(C)** Marker gene expression and TF motif scores for each cluster. **(D)** Heatmap showing the proportion of cells in the RNA cluster that overlaps with chromatin-defined and methylation-defined clusters. **(E)** UMAP colored by the percentage of control cells in neighboring cells. **(F)** DNA methylation was depleted upon *Uhrf1* knockout. **(G)** Volcano plot of differentially expressed genes between *Uhrf1* knockout and wild-type identified from neural crest cells. **(H)** Gene Ontology (GO) term enrichment in differentially expressed genes identified from neural crest cells.

Leveraging ME-seq, we assessed the congruence between the epigenome and transcriptome across an atlas of 24,726 high-quality single-cell profiles (**Extended Data Fig.2A**). This dataset represents a unique trimodal single-cell mouse embryo atlas, enabling comprehensive analysis of the transcriptomic and epigenomic alterations associated with UHRF1 loss. We next evaluated several technical quality metrics. First, RNA counts, ATAC counts, and DNA methylation levels were highly correlated across technical replicates from the same sample (average Pearson correlation = 0.99), demonstrating high consistency in our sample preparation. Second, doublets and cells with elevated mitochondrial gene expression accounted for only a small fraction of total cells. Third, we assessed the 5mC conversion efficiency using fully methylated control DNA and achieved over 90% conversion efficiency. Fourth, we confirmed the deletion of exons 6–10 in the UHRF1 knockout embryos (**Extended Data Fig.2B**).

To define cell subsets, we clustered the RNA modality of the ME-seq data (**Fig.2B**). The cell types are annotated based on canonical marker genes (**Fig.2C**) previously described in the literature^26^. We captured all the major expected cell types in the embryos, including ectoderm lineage (e.g., forebrain/midbrain, hindbrain, neural crest), mesodermal lineage (e.g., allantois, amniochorionic mesoderm, endothelium, and first heart field), and endodermal lineage (e.g., gut and extraembryonic visceral endoderm) (**Fig.2C**). We then projected the cell type annotation onto the ATAC and DNA methylation modalities of the ME-seq data and found that both projections recapitulated the RNA-based clustering (**Fig.2B**). ME-seq not only resolved major germ layers but could also distinguish closely related subtypes (e.g., first vs. second heart field). Moreover, cell memberships inferred from scATAC-seq and scDNAme-seq clusters were highly congruent with those from scRNA-seq clusters (**Fig.2D**), consistently identifying the same major cell types. Importantly, these RNA-based clusters showed distinct chromatin landscapes in TF motif scores (**Extended Data Fig.2C**) and characteristic marker gene accessibility (**Extended Data Fig.2D**) derived from scATAC-seq data.

Next, we sought to identify the cell state differences caused by the knockout. Upon knockout, all cell types were conserved (**Extended Data Fig.2E**) but displayed subtle shifts in relative abundance (**Fig.2E, Extended Data Fig.2F**). Specifically, we observed significantly reduced proportions of gut, paraxial mesoderm, and forebrain/midbrain populations, accompanied by increased abundances of amniochorionic mesoderm and extraembryonic visceral endoderm populations. Furthermore, we assessed changes in the global methylation level. As expected, UHRF1 knockout dramatically reduced methylation levels in the regulatory regions defined by scATAC-seq (*p* < 2.2×10^-16^, *t*-test, **Fig.2F**) as well as across gene bodies (**Extended Data Fig.2G**).

### UHRF1 knockout is associated with alterations in CTCF binding

As the *Uhrf1* knockout caused pronounced defects in the neural crest complex, we further examined the accompanying transcriptomic alterations. In the neural crest population, we identified 150 high-confidence differentially expressed genes (DEGs; p < 0.01, fold change > 1, Wilcoxon rank-sum test, **Fig.2G**). These DEGs were strongly enriched in functional terms, including *Wnt* signaling pathway, neuron differentiation, and chromatin organization and remodeling (**Fig.2H)**, suggesting that UHRF1 disruption may be directly associated with chromatin changes. To test this possibility, we next examined the chromatin alterations in the neural crest and other cell populations.

Upon UHRF1 knockout, we identified 3,090 upregulated and 4,232 downregulated differential peaks (adjusted *p* < 0.1, fold change > 2, logistic regression; **Fig.3A**). These differential peaks were strongly enriched for the CTCF binding motif, as well as MYF5 and NFIX motifs (**Fig.3B**). CTCF is known to play a critical role in organizing the three-dimensional genome architecture. Acting as both an insulator and an architectural protein, CTCF influences gene expression and chromatin organization by binding to DNA and interacting with other proteins, such as cohesin. These interactions enable CTCF to form chromatin loops and establish boundaries within the genome, creating TADs. TADs are essential for regulating gene expression by controlling interactions between enhancers, promoters, and other regulatory elements^27^.

**Figure 3.**
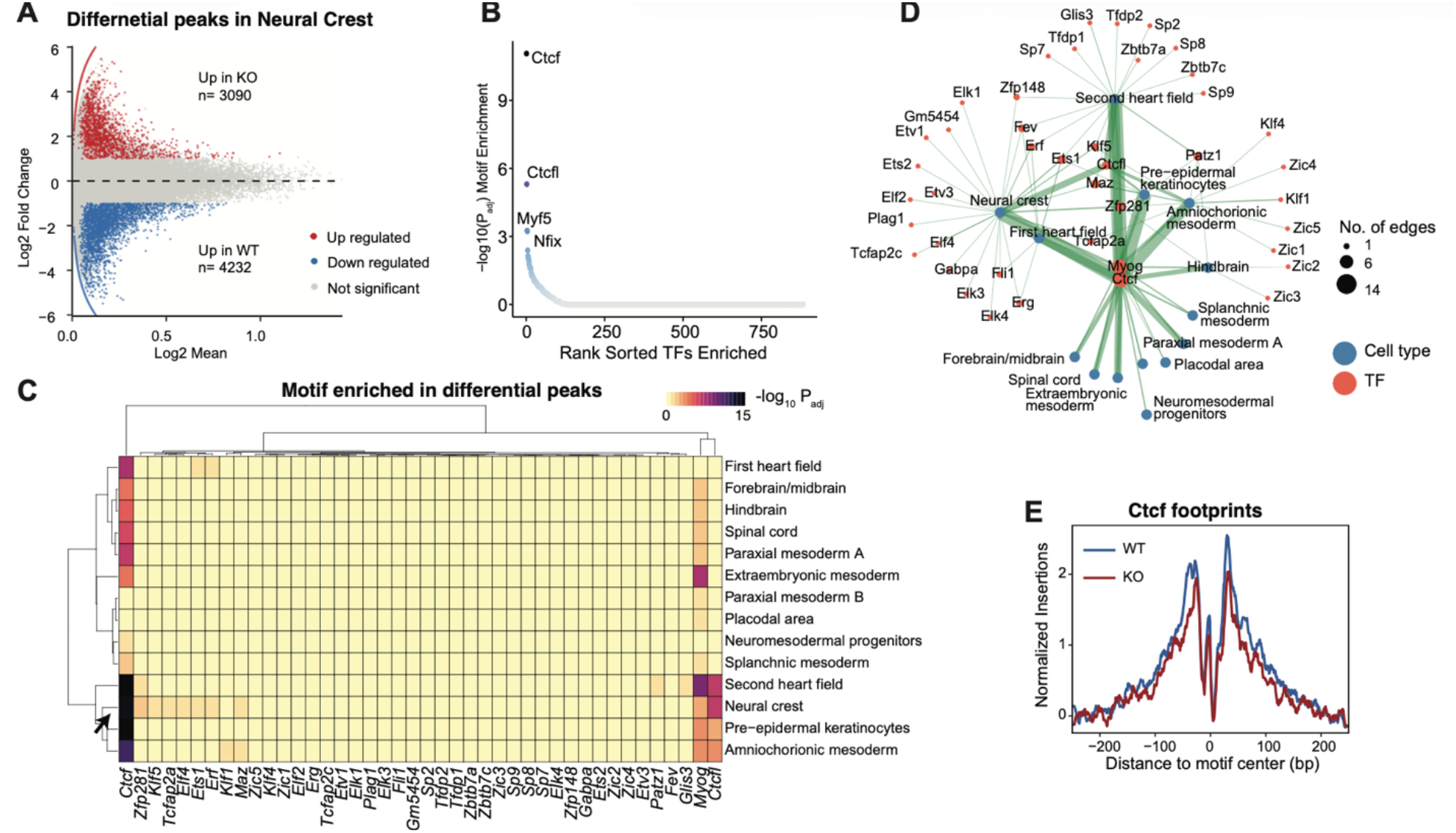
*Uhrf1* knockout is associated with alterations in CTCF binding. **(A)** MA plot of differential accessible peaks between *Uhrf1* knockout and wild-type identified from neural crest cells. **(B)** Ranked TF motifs enriched in the neural crest differential peaks. **(C)** TF motif enrichment in differential peaks between *Uhrf1* knockout and wild-type embryos across all cell types. **(D)** TF regulatory network showing the driver TF for each cell type. The width of an edge indicates the significance of enrichment (−log_10_(FDR)). **(E)** CTCF footprints in neural crest cells.

Importantly, the CTCF motifs were not only enriched in differential peaks in neural crest, but also in many other, though not all, populations, such as amniochorionic mesoderm (adjusted *p* < 10^-15^), whose abundance was significantly changed upon knockout (**Fig.3C)**. Furthermore, network analysis placed CTCF as the central hub (**Fig.3D)**, directly regulating neural crest and other populations. Consistent with this, TF footprint analysis confirmed reduced CTCF binding in the neural crest (**Fig.3E**). Interestingly, these changes in CTCF binding were not accompanied by alterations in *Ctcf* expression or chromatin accessibility at the *Ctcf* locus (**Extended Data Fig. 3A**), suggesting that reduced CTCF occupancy is not driven by transcriptional downregulation. To determine whether CTCF-associated regulatory regions were affected by the global methylation loss caused by UHRF1 deficiency, we quantified CpG methylation within chromatin-accessible peaks harboring CTCF motifs. *Uhrf1* knockout embryos exhibited significantly reduced methylation at these sites compared with wild-type embryos (**Extended Data Fig. 3B**), consistent with the broad hypomethylation observed genome-wide. We further performed CTCF CUT&RUN and examined DNA methylation levels within called CTCF peak regions. Methylation levels at these CTCF-associated regions were similarly reduced in *Uhrf1* knockout embryos relative to wild-type embryos (**Extended Data Fig. 3C**). Because methylation loss was widespread across the genome, these data do not establish that CTCF sites are preferentially demethylated. Rather, they show that CTCF-associated regulatory regions are included among the genomic regions affected by UHRF1-dependent methylation loss, providing a chromatin context for the altered CTCF-associated regulatory architecture observed in knockout embryos.

Although CTCF RNA expression and chromatin accessibility at the CTCF locus were largely unchanged upon UHRF1 loss, these measurements do not exclude altered CTCF regulation at the protein or chromatin-association level. We therefore examined bulk protein abundance by mass spectrometry-based proteomics. Only a small number of proteins were significantly altered in UHRF1 knockout embryos, including the expected reduction of UHRF1, whereas total CTCF protein abundance was not significantly changed (*p* > 0.05, n = 3 per group). These results indicate that UHRF1 loss does not broadly reduce CTCF expression or protein levels at the whole-embryo level (**Extended Data Fig. 3D**). Because UHRF1 can regulate chromatin through mechanisms beyond DNA methylation maintenance, we next examined whether UHRF1 loss alters histone modification landscapes. H3K27me3 CUT&RUN identified 1,115 differential peaks between wild-type and UHRF1-null embryos, indicating altered Polycomb-associated chromatin states upon UHRF1 loss. Of these differential H3K27me3 peaks, 859 overlapped ATAC peaks, and 692 of the 859 ATAC-overlapping peaks occurred at ATAC peaks containing CTCF motifs.

Compared with the frequency of CTCF motif-containing peaks among all ATAC peaks, this represented a significant enrichment (80.6% versus 29.3%; 2.75-fold enrichment; hypergeometric test, *p* ≈ 7.85 × 10⁻²¹³). These results suggest that H3K27me3 alterations converge with a subset of accessible CTCF-associated regulatory regions.

### UHRF1 alters BMP signaling in the neural crest and cell-cell interactions

We reasoned that the chromatin alterations caused by the UHRF1 knockout may influence beyond neural crest cells by disrupting cell-cell interactions (CCIs). To test this hypothesis, we carried out cell-cell communication analysis on wild-type and knockout embryos, respectively. Upon knockout, we observed a global reduction in CCIs (from 33,757 to 26,581) and across nearly all cell types except for the extraembryonic visceral endoderm (**Fig.4A,B, Extended Data Fig.4A**). In particular, the knockout led to dramatic decreases in neural crest-related cell-cell interactions (**Fig.4C,D, Extended Data Fig.4B**). Importantly, comparable cell numbers and data quality were recovered from wild-type and knockout samples, ruling out potential confounding effects.

**Figure 4.**
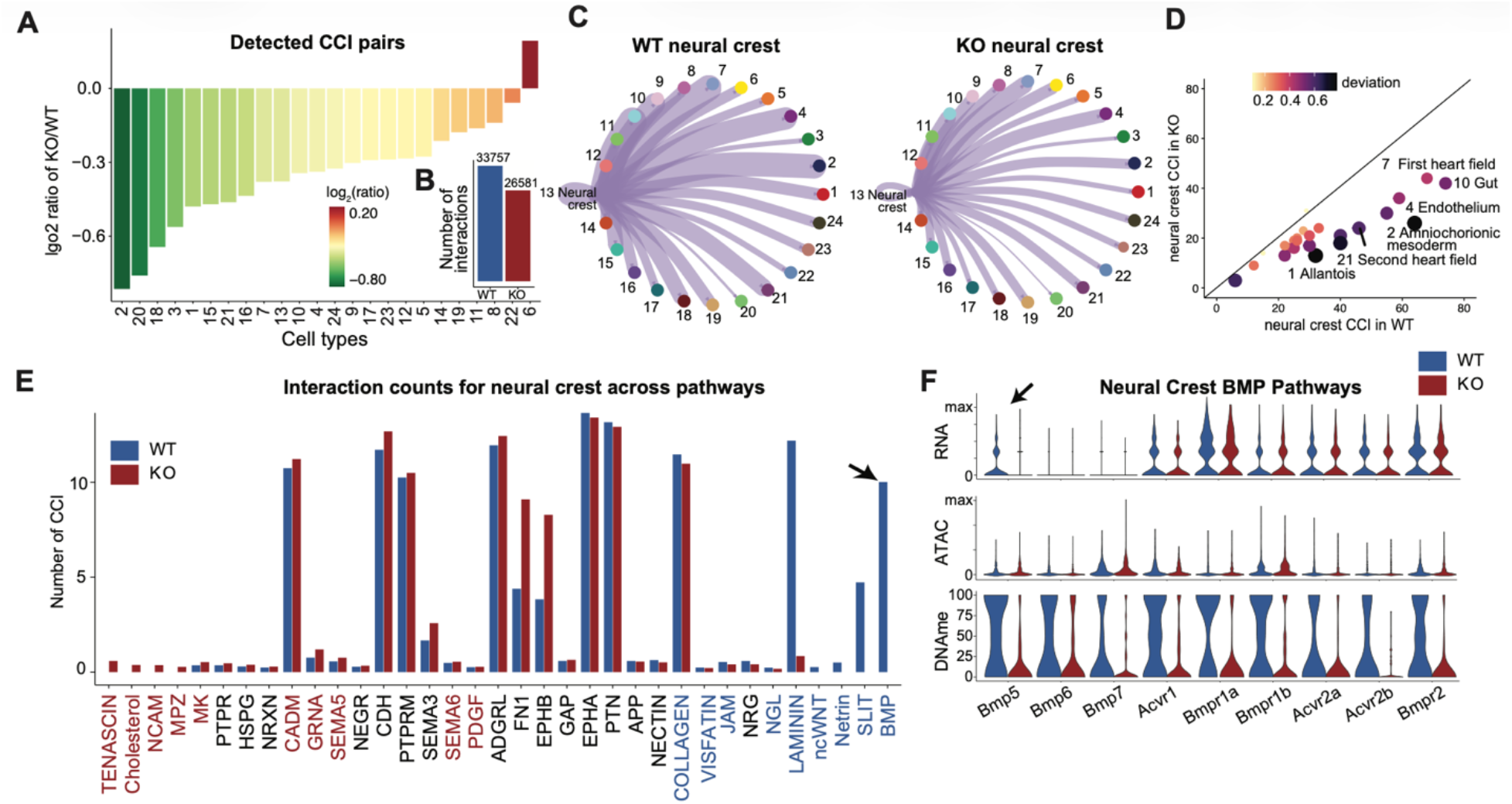
*Uhrf1* alters predicted BMP signaling in neural crest and cell-cell interactions. (A-B) The number of cell-cell interactions was significantly reduced by *Uhrf1* knockout in each cell type (A) and globally (B). **(C-D)** Cmparing the cell-cell interactions from neural crest to other populations. **(E)** Examining the number of neural crest cell-cell interactions in each pathway, including BMP signaling. **(F)** Comparing the RNA expression (top), ATAC (middle) and DNA methylation (bottom) signals around known ligand-receptor genes in BMP signaling pathway. ATAC is defined using gene score in ArchR (promoter and gene body). DNA methylation is defined as average gene body methylation.

Next, we examined the signaling pathways identified in the neural crest. Strikingly, the BMP pathway activity was predicted to be largely reduced based on ligand–receptor analysis in the knockout (**Fig.4E**). The BMP signaling pathway, a major branch of the TGF-β superfamily, regulates diverse cellular processes including growth, apoptosis, and differentiation, and plays an essential role in both development and tissue homeostasis^28^. We further analyzed all known BMP signaling components and found that only *Bmp5* showed significant expression changes in the neural crest (**Fig.4F**). Although no significant differences in promoter or gene-body chromatin accessibility were observed for *Bmp5* or other BMP pathway genes, all exhibited consistent reductions in DNA methylation (**Fig.4F**). These results suggest that UHRF1 loss–induced hypomethylation does not primarily affect proximal regulatory elements but may instead alter distal *cis*-regulatory regions linked to these genes.

### *Cis*-regulation analysis identified chromatin interaction changes

To dissect these, we applied the *cis-regulation* analytical framework we previously developed^29^ to identify peak–gene associations, based on the co-variation of chromatin accessibility and gene expression across cells, while controlling for technical biases in chromatin accessibility measurements. Further, we extended this framework to identify methylation–gene associations based on the co-variation in DNA methylation and gene expression across cells (**Fig.5A**). We identified several genes with an exceptionally large (>18) number of significant peak-gene associations (FDR < 0.05, *z*-test), such as *Ptch1* and *Slit3* (**Fig.5C**). *Ptch1* is a key component of the SHH signaling pathway, which is essential for proper cell differentiation, tissue polarity, and cell proliferation during embryonic development^30^. *Slit3* promotes developmental angiogenesis to support organogenesis during mammalian development^31^. We also identified several genes with an exceptionally large (> 6) number of significant methylation-gene associations (*FDR* < 0.05, *z*-test), such as *Hoxc4*, *Ptch1*, and *Cldn6* (**Fig.5B**). *Hoxc4* is known for regulating organogenesis and cell differentiation. We identified a similar number of genes with an exceptionally large (>6) number of significant peak-gene associations and significant methylation-gene associations (**Extended Data Fig.5A**) in the knockout. However, a subset of genes showed significant changes in the number of associated peaks or methylation sites (**Extended Data Fig.5B**). For example, *Peg3*, a gene that plays a critical role in cell proliferation and p53-mediated apoptosis^32^, was found to have five additional significant peak–gene associations and seven additional significant methylation–gene associations.

**Figure 5.**
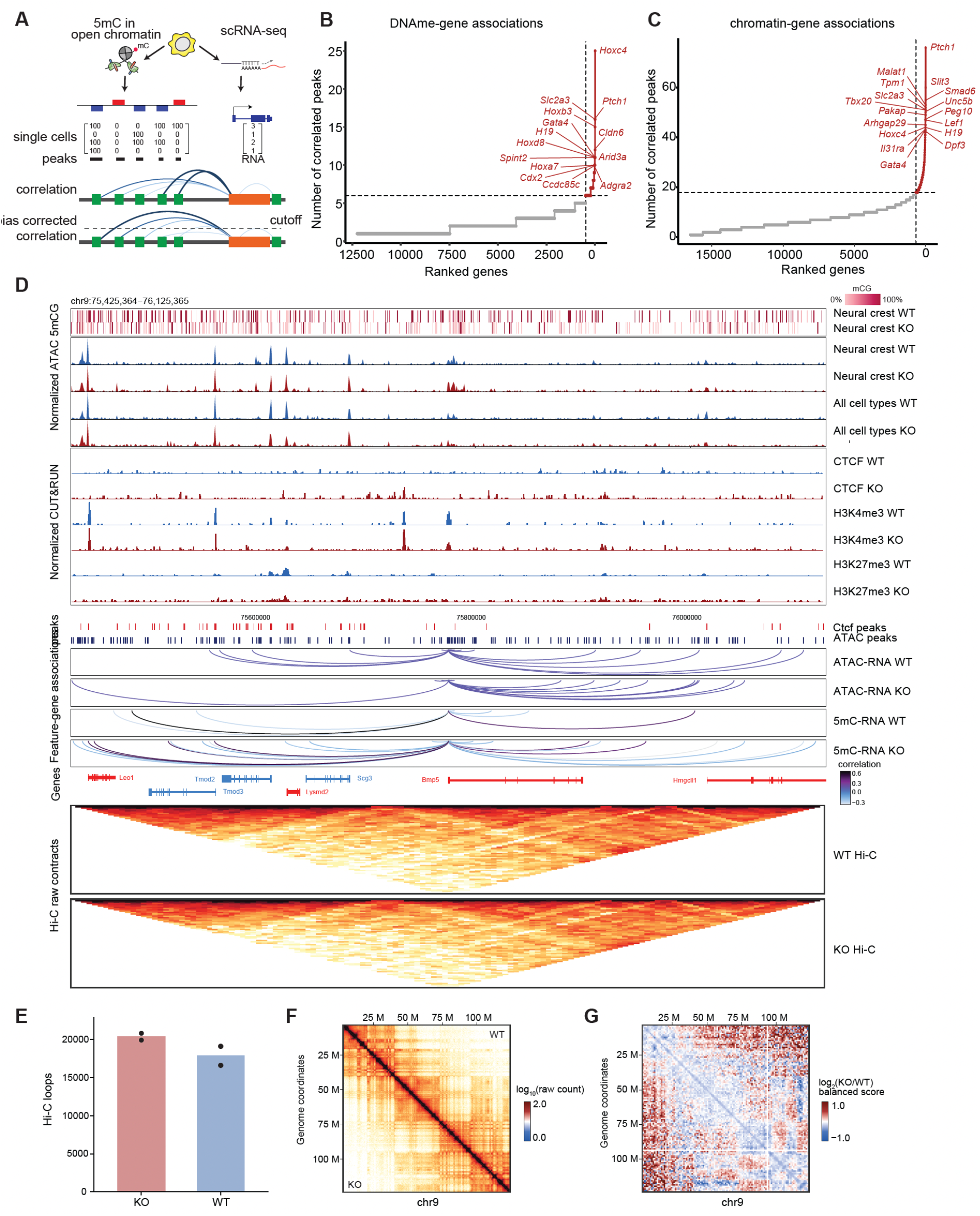
*Cis*-regulation analysis identifies chromatin and 3D genome alterations associated with UHRF1 loss. **(A)** Schematic depicting an analytical framework for analysis of distal regulatory elements and expression of genes. **(B-C)** The number of significantly correlated peaks (p < 0.05) for each gene (50 kb from TSSs). The correlation to RNA expression was calculated using DNA methylation (B) or accessibility (C) in peaks. **(D)** Genome browser view of the *Bmp5* locus showing DNA methylation, normalized ATAC signal, CUT&RUN, feature-gene associations, and Hi-C contact maps in wild-type and knockout embryos. Loops denote correlations between regulatory features and Bmp5 RNA expression, with loop height representing the significance of the correlation and color indicating correlation strength. **(E)** Number of Hi-C loops detected in wild-type and knockout embryos. **(F)** Representative Hi-C contact map on chr9 showing chromatin contact patterns in wild-type and knockout embryos. **(G)** Differential Hi-C contact map showing the log_2_ ratio of knockout versus wild-type balanced contact scores on chr9.

Cell–cell communication analysis revealed reduced *Bmp5*-related ligand–receptor interactions in UHRF1-null embryos, particularly in neural crest cells. This reduction was accompanied by significantly decreased *Bmp5* expression and altered regulatory features at the *Bmp5* locus. Focusing on the *Bmp5* gene, we found a similar number of peak–gene associations (19 and 18 for wild-type and knockout, respectively) but a dramatically increased number of methylation-gene associations (10 and 27 for wild-type and knockout, respectively). Likewise, we found more methylation-peak associations in *Hoxc4* and many other genes (**Extended Data Fig.5C**). Although global DNA hypomethylation is often associated with de-repression of regulatory elements or genes, its transcriptional consequences are highly dependent on genomic context^33^. At the *Bmp5* locus, UHRF1 loss was associated with decreased H3K4me3 signal at the *Bmp5* promoter, indicating reduced promoter activity. Thus, *Bmp5* downregulation is unlikely to reflect a simple direct relationship between DNA hypomethylation and transcriptional activation. Instead, these findings suggest that UHRF1 loss can disrupt locus-specific chromatin states required for gene activation, potentially through altered promoter activity, enhancer connectivity, or lineage-specific regulatory programs.

To explore why neural crest cells are particularly vulnerable to UHRF1 loss, we compared regulatory architecture across embryonic lineages. Neural crest cells exhibited relatively low promoter-proximal accessibility, as indicated by reduced TSS enrichment, and showed fewer detectable ATAC–RNA peak–gene associations compared with many other lineages (**Extended Data Fig.6A,B**). Because these measurements can be influenced by chromatin data quality, we interpret this pattern cautiously. Nonetheless, together with the pronounced transcriptional, chromatin, and cell–cell communication changes observed in neural crest cells, these findings suggest that neural crest identity may depend on a regulatory program that is particularly sensitive to disruption of DNA methylation-dependent chromatin organization. Thus, UHRF1 loss may impair neural crest development not only through global hypomethylation, but also by weakening the enhancer–gene and chromatin interaction landscape required for lineage stabilization. Future lineage-resolved chromatin profiling will be needed to determine whether neural crest cells possess uniquely vulnerable CTCF-or enhancer-dependent regulatory architecture.

### UHRF1 loss disrupts CTCF-associated chromatin occupancy and 3D genome organization

The enrichment of CTCF motifs among differentially accessible peaks and the reduction in ATAC-derived CTCF footprinting suggested that UHRF1 loss may perturb CTCF-associated regulatory architecture. However, these analyses infer CTCF activity from chromatin accessibility and motif structure rather than directly measuring CTCF chromatin occupancy at chromatin. We therefore performed CTCF CUT&RUN in wild-type and UHRF1-null embryos. Consistent with the ME-seq-based inference, UHRF1-null embryos showed reduced CTCF CUT&RUN library complexity and decreased CTCF-associated chromatin signal at CTCF peak regions, indicating impaired CTCF chromatin association upon UHRF1 loss (**Extended Data Fig.6C,D**). Together, these findings support a model in which UHRF1 loss perturbs CTCF-associated chromatin organization, potentially through altered chromatin occupancy or stability rather than reduced CTCF expression.

To directly examine whether these CTCF-associated changes were linked to altered higher-order chromatin organization, we performed Hi-C in wild-type and UHRF1-null embryos. Hi-C contact maps were reproducible across biological replicates and revealed broad changes in chromatin contact organization upon UHRF1 loss (**Fig.5F,G; Extended Data Fig.6E**). Genome-wide, UHRF1-null embryos showed reduced *cis*-contact frequency, indicating weakened intra-chromosomal contact organization. Interestingly, despite this reduction in overall *cis* contacts, the number of called chromatin loops increased in knockout embryos (**Fig.5E**). These observations are not mutually exclusive, as loop calling identifies focal interactions enriched relative to local background rather than total chromatin connectivity. Thus, reduced broad *cis*-contact frequency in UHRF1-null embryos may increase the relative prominence of a subset of focal interactions, reflecting redistribution of chromatin contacts, potentially through altered CTCF/cohesin occupancy at normally methylated regulatory sites.

Together, these analyses extend the ME-seq findings by showing that UHRF1 loss is accompanied by reduced CTCF chromatin association and broad remodeling of chromatin contacts. These results support a model in which UHRF1-dependent methylation maintenance acts together with histone-state regulation and architectural proteins to preserve lineage-specific regulatory programs during early embryogenesis.

## Discussion

Our study demonstrates that murine UHRF1 plays an essential role in maintaining DNA methylation and chromatin architecture during early mouse embryonic development. Using a knockout system in ESCs and a novel multi-modal single-cell approach (ME-seq) in embryos, we show that while UHRF1 is dispensable for pluripotency and teratoma formation *in vitro*, it is indispensable *in vivo* for embryogenesis beyond E8.5. Loss of UHRF1 leads to global DNA hypomethylation, widespread chromatin dysregulation, and ultimately embryonic lethality. UHRF1 is known to directly interact with DNMT1 in order to maintain methylation patterns across DNA strands at CpG sites during DNA replication^34^, and loss of DNMT1 similarly causes global DNA hypomethylation and embryonic lethality^17^. These findings underscore that the maintenance of DNA methylation is essential for post-implantation development.

Our *in vitro* experiments demonstrate that UHRF1 deficiency leads to near-complete depletion of 5mC and 5hmC without affecting ESC self-renewal. This observation aligns with previous studies showing that fast-proliferating ESCs can tolerate global hypomethylation^35^, likely due to redundant mechanisms maintaining core pluripotency transcriptional networks. However, this tolerance does not extend to the complex demands of embryonic development, as reflected by the severe defects and lethality we observed at E9.5.

Moreover, UHRF1-null ESCs showed modest transcriptional changes, including bidirectional changes of several imprinted genes. The up-regulation of Xlr3/4 reflects the loss of constitutive gene-body/repeat methylation that maintains repression at this rodent-specific, X-linked locus, rather than the erasure of a parent-of-origin imprint. The *Xlr* brain imprint has no classic DMR (both alleles are methylated; paternal silencing is enforced post-transcriptionally), and its applicability to ESCs is unresolved. This places the *Xlr* signal alongside the broader de-repression of germline genes, retrotransposons, and X-linked repeat families (*Xlr*/*Xmr*, *Rhox*, *MAGE*) characteristic of methylation-deficient ESCs. Although these changes were not sufficient to disrupt pluripotency or teratoma formation *in vitro*, they may indicate loss of epigenetic dosage control at methylation-sensitive loci. During embryogenesis, such imprinting defects could contribute to developmental instability by altering the dosage of genes involved in growth, signaling, or lineage specification. However, the selective vulnerability of neural crest cells is unlikely to be explained by imprinted gene deregulation alone. Instead, our *in vivo* analyses suggest that imprinting changes occur alongside broader defects in chromatin state, CTCF-associated organization, *cis*-contact architecture, and developmental signaling programs. We therefore interpret imprinted gene upregulation as part of the broader UHRF1-dependent epigenomic disruption that may sensitize embryonic lineages to failure during post-implantation development.

Through ME-seq, which simultaneously captures the transcriptome, chromatin accessibility, and DNA methylation in single cells, we created a trimodal atlas of wild-type and UHRF1-null embryos. These data reveal that UHRF1 loss causes global hypomethylation across regulatory regions and gene bodies, accompanied by altered chromatin accessibility and substantial shifts in lineage composition. Notably, we observed reductions in forebrain/midbrain, gut, and paraxial mesoderm populations, and expansion of extraembryonic lineages. These findings highlight that DNA methylation fidelity, mediated by UHRF1, is essential for appropriate lineage allocation.

Our analysis identifies CTCF as a central factor whose function is altered in the absence of UHRF1. Disruption of CpG methylation alters genomic CTCF occupancy, likely perturbing enhancer–promoter topologies consistent with the observed defects in specific cell-lineage populations resulting from altered transcriptional patterns. We observed significant differences in chromatin peaks enriched for CTCF motifs, with TF footprinting confirming altered CTCF occupancy.

Another major consequence of UHRF1 loss was a reduction in intercellular communication, particularly among neural crest lineages, and a predicted reduction in BMP-related communication, an essential pathway for neural crest differentiation and tissue patterning. While *Bmp5*^28^ was the only BMP family gene to show significant transcriptional downregulation, our *cis*-regulatory analysis revealed an increase in distal methylation-gene associations rather than promoter accessibility changes, implicating long-range enhancer rewiring as a key mechanism of dysregulation. Genes such as *Ptch1*, *Slit3*, and *Hoxc4*—critical mediators of developmental signaling—were particularly affected, highlighting a broad impact on morphogenetic signaling networks.

Because UHRF1 is a multifunctional chromatin regulator that recognizes both methylated DNA and histone modification states, the regulatory defects observed in UHRF1-null embryos are unlikely to reflect DNA methylation loss alone. Consistent with this broader role, CUT&RUN profiling revealed altered histone modification landscapes together with changes in CTCF-associated chromatin organization. Thus, UHRF1 loss appears to perturb an integrated chromatin regulatory axis involving DNA methylation maintenance, histone modification states, and CTCF-linked *cis*-regulatory architecture. These combined defects may contribute to impaired enhancer activity, altered peak–gene associations, and failed lineage stabilization during post-implantation development.

Because UHRF1-null mESCs retained pluripotency and did not exhibit overt *in vitro* defects, we focused our CTCF CUT&RUN and Hi-C analyses on embryos, in which UHRF1 loss produced clear developmental and lineage-specific phenotypes. The CTCF-associated chromatin changes observed *in vivo* may therefore reflect both direct effects of UHRF1-dependent methylation loss and developmental context-specific regulatory changes that emerge during lineage specification. Because neural crest cells represent only a small fraction of the E8.5 embryo^36^, whole-embryo Hi-C, CUT&RUN, and proteomic analyses are informative for defining global chromatin and protein-level changes but cannot directly resolve whether these alterations occur specifically within neural crest cells. Future studies using mESC differentiation models, including directed neural crest differentiation, will help define which CTCF changes are present in pluripotent cells and which arise during lineage commitment.

UHRF1-null embryos showed reduced *cis*-contact frequency and altered chromatin loop organization, indicating broad disruption of 3D genome architecture. Such disruption may affect the regulatory environment of developmental signaling genes. Consistent with this possibility, Bmp5 expression was reduced in knockout embryos and was accompanied by decreased H3K4me3 at the Bmp5 promoter and altered local regulatory organization. Together, these findings suggest that reduced predicted BMP-related ligand–receptor interactions may arise, at least in part, from impaired chromatin regulation of signaling pathway components during neural crest development.

Overall, these findings support a model in which UHRF1-dependent methylation maintenance acts together with histone-state regulation and CTCF-associated chromatin organization to preserve chromatin contact architecture and developmental gene regulation during early embryogenesis. Loss of UHRF1 therefore disrupts multiple interconnected regulatory layers, including DNA methylation, histone modification landscapes, CTCF-associated chromatin signal, chromatin contacts, and predicted developmental signaling communication. These combined defects may impair lineage stabilization, with neural crest cells emerging as a particularly vulnerable population.

### Limitations of the study

These results advance our understanding of how DNA methylation and 3D genome organization cooperate during early development, but they also raise several important questions. How does hypomethylation specifically impair CTCF binding in some loci but not others? Could the observed phenotypes be partially rescued by stabilizing CTCF-cohesin loops? Because we did not perform rescue experiments with wild-type or separation-of-function UHRF1 mutants, the current study cannot distinguish the relative contributions of UHRF1’s DNA methylation maintenance function from its histone-and chromatin-binding functions. Future rescue studies will be required to determine which molecular activities of UHRF1 are necessary for maintaining enhancer activity, CTCF-associated chromatin organization, and embryonic lineage stability. A limitation of the current study is that we did not directly measure CTCF nuclear localization, protein stability, or cohesin occupancy. Because CTCF-mediated loop formation depends on cohesin, future studies examining RAD21, SMC1A, SMC3, and related cohesin regulators in lineage-resolved embryo material will be important for defining how UHRF1 loss disrupts CTCF/cohesin-dependent genome organization. In addition, our Hi-C, CUT&RUN, and proteomic analyses were performed using whole-embryo material. Because neural crest cells comprise only a small subset of the E8.5 embryo, these assays therefore provide direct evidence for global changes in chromatin contact organization, CTCF-associated chromatin signal, histone modification landscapes, and protein abundance, but they do not resolve these changes in a neural crest-specific manner. Future lineage-resolved chromatin profiling will be needed to determine the extent to which the altered chromatin architecture observed at the whole-embryo level occurs specifically within neural crest cells. Another limitation of the current study is that the genome-wide nature of DNA hypomethylation in UHRF1 knockout embryos prevents assignment of specific chromatin interaction changes to local methylation loss alone. Future targeted methylation editing or separation-of-function rescue experiments will be required to distinguish methylation-dependent effects from other chromatin regulatory functions of UHRF1. Future profiling of additional UHRF1-linked histone marks, including H3K9me2/3 and histone ubiquitination, will help define how histone-state regulation cooperates with DNA methylation maintenance to preserve chromatin architecture and lineage stability. Additionally, future studies could explore whether similar mechanisms underlie developmental disorders and cancers in which UHRF1 dysfunction perturbs DNA methylation and chromatin organization.

**Supplemental Table 1. Imprinted genes differentially expressed in Uhrf1 KO versus WT ESCs.** Significant imprinted loci (adjusted *p* < 0.10, fold change > 1.5) from RNA-seq of Uhrf1 KO and WT ESCs, ranked by log₂ fold-change (KO/WT). For each gene, the parent-of-origin annotation, log₂ fold-change, Benjamini–Hochberg-adjusted *p*-value (FDR), direction of change, and mean normalized read counts (n = 3 per genotype) are shown. Loci up-regulated in KO are shaded orange; those down-regulated are shaded blue. The response is bidirectional: of 17 significant imprinted genes, 13 are up-regulated and 4 are down-regulated. Loss of DNA methylation derepresses genes that methylation normally silences (i.e., *Xlr3b*) while reducing expression of genes dependent on an intact methylation imprint (i.e., *Cdkn1c*, *Grb10*).

**Supplemental Table 2. Embryo recovery rates of UHRF1 mouse crosses. Resource availability**

**Lead contact.**

Sai Ma

## Materials availability

All unique/stable reagents generated in this study are available from the Lead Contact with a completed Materials Transfer Agreement.

## Acknowledgments

We thank all the members of the Walsh and Ma labs for critical reading of the manuscript and helpful discussions. S.M acknowledges support from the NIH through awards U01CA290442, R61CA297881, and R21CA301237, and Mount Sinai FBI scholar. M. J. W. acknowledges support from NIH through awards R01GM119189, R01DK116873, and R01DK118946.

## Author contributions

A.K. generated the mESC and mouse models for constitutive *Uhrf1+/-* deletion, maintained the mouse colonies, performed staining and functional validation, collected mouse embryos, with help from L.H., and generated bulk sequencing data and analysis. B.Z. generated the embryonic single-cell sequencing library and computational analysis with participation of A.K and Y.S. S.M provided computational support. Y.S. performed methyl-seq bulk sequencing analysis. A.C. performed cell-cell communication analysis. N.J. maintained initial Uhrf1*^flox/flox^* mouse colonies, their genotypes and ESC colonies derived from embryos. Y.S. and P.S. performed proteomics experiment and analysis. S.M., B.Z., A.K. and Y.S. conducted the data analysis with direct input from M.J.W. All authors participated in writing the manuscript. M.J.W. and S.M. conceived and supervised the research, and all authors reviewed the manuscript.

## Material and Methods

### Cell culture and manipulation

Embryonic stem cells (ESCs) were derived from *Uhrf1*^Fx/Fx^ 129/Sv:C57BL/6-mice and cultured in 2i/LIF as described^18,37^. *Uhrf1*^-/-^ ESCs were obtained by transfecting *Uhrf1*^Fx/Fx^ ESCs with a linearized CreERT2-IRES-Blasticidin plasmid^38^ using Lipofectamine 2000. ESCs were cultured for 48 h in 2i/LIF containing 1 μg/ml 4-hydroxytamoxifen (4-OHT) to induce CRE recombination, then selected for 48 h with 1 μg/ml blasticidin to retain cells expressing the CreERT2-IRES-blasticidin cassette. Resistant clones were picked and expanded, and UHRF1-null status was confirmed by genotyping and Western blot. Cells were cultured for about two weeks before analysis. Cell cycle phases were determined using the Muse Cell Cycle Assay Kit (Merck Millipore) and analyzed on the Muse instrument.

### Embryo dissection and culture

*Uhrf1*^+/−^ male and female mice were naturally mated and checked every morning for the appearance of a vaginal plug on E0.5. E3.5 embryos were flushed with PBS/FBS from the uterus and E4.5 and older embryos were dissected from the uterus in PBS/FBS and fixed in 4% PFA for 15 min to overnight according to size. The placenta was used for genotyping. Forward primer GATAAGCTTACATAATCACATGGA and reverse primer CCTATGAAATCCTTTGCTGC were used to determine the sex. LNL5F ATTTGGCTTTCTGGATGCTTCTGTT, FNFL5F TTGATGTTCACAGGTCAGTGAGTCC, and FNL3R TGTGACTTCCCATCTTCCCTTCTAA were used to distinguish between WT and Het *Uhrf1* embryos. 2x PCR master mix (Promega) was used and 7% DMSO was added for the PCR amplification with annealing temperature at 59°C.

### Mice

*Uhrf1*^Fx/Fx^ mice were obtained on a J1 background. *Uhrf1*^-/+^ alleles were generated from the floxed alleles by crossing with B6.C-Tg(CMV-cre)1Cgn/J (Jackson Laboratory) and subsequently crossing out the Cre driver. All animal experimentation was conducted according to protocols approved by the Institutional Animal Care and Use Committee of Icahn School of Medicine at Mount Sinai.

### Genomic DNA Dot-blot for mC/hmC

Genomic DNA was isolated using the DNeasy Blood & Tissue Kits (Qiagen). 20 μl denaturing buffer (0.8 M NaOH, 200 mM EDTA) was added to 3 μg DNA in 20 μl H2O and incubated at 95°C for 10 min. 20 μl of cold 2 M ammonium acetate solution was added for neutralization and mixed well. Two serial dilutions (1:2) and (1:4) were prepared. 4 μl (200 ng, 100 ng, 50 ng) of each dilution was loaded at each dot on the nitrocellulose membrane. The membrane was air-dried in a 65 °C oven for 30 min. DNA was auto-crosslinked with 1200×100 uJ/cm^2^. After a rinse with TBST the membrane was blocked with 5% milk/TBST for 1 hour at RT. The membrane was incubated at RT 1-2 h using the following antibodies: anti-mC (Millipore MABE164, 33D3, 1:1000, Ms) and anti-hmC (active motif 39769, 1:5000∼10000). Membrane was washed 3 times for 10 minutes with TBST and visualized using ECL detection (Thermo Scientific).

### Immunoblotting

Nuclear extracts were resolved by SDS-PAGE, transferred to nitrocellulose membranes (GE Healthcare Life sciences), and immunoblotted with the indicated with *Uhrf1* antibody (Santa Cruz, sc-373750) followed by ECL detection (Thermo Scientific).

### Immunofluorescence staining

Immunostaining was performed as described^39^. Embryos or ESCs were fixed 15 minutes in 4% PFA, permeabilized for 30 minutes in 0.5% Triton X100 (T9284, Sigma) and 3 mg/ml PVP in PBS, blocked in 0.1% BSA with 2% donkey serum in 0.25% Triton X100 in PBS/PVP for a minimum of 30 minutes and incubated overnight with primary antibodies used at 1/200 dilution. Primary antibodies used: NANOG (Cosmo Bio Co., Ltd, RCAB0002P-F), E-Cadherin (R&D Systems, AF1700), and Cl-Casp3 (Cell Signaling, 9661). On the next day, embryos were washed in PBST and incubated for 1 hour with secondary Alexa Fluor (Invitrogen) conjugated antibodies diluted 1/500 in 1% donkey serum in PBST. DAPI was used to counterstain and Vectashield to mount the samples. Confocal images were taken using a Leica TCS SP5 confocal microscope. LAS AF was used to visualize confocal images.

### RT-qPCR analysis

RNA was extracted using the RNeasy Mini Kit (Qiagen). Up to 1 μg total RNA was reverse transcribed using the iScript™ Reverse Transcription Supermix (Bio Rad). Quantitative PCR was performed using the Luna Universal qPCR Master Mix (NEB) on the Stratagene Mx3005P Real-Time PCR System (Agilent Technologies). Gene expression-specific primers used for this study are listed below.

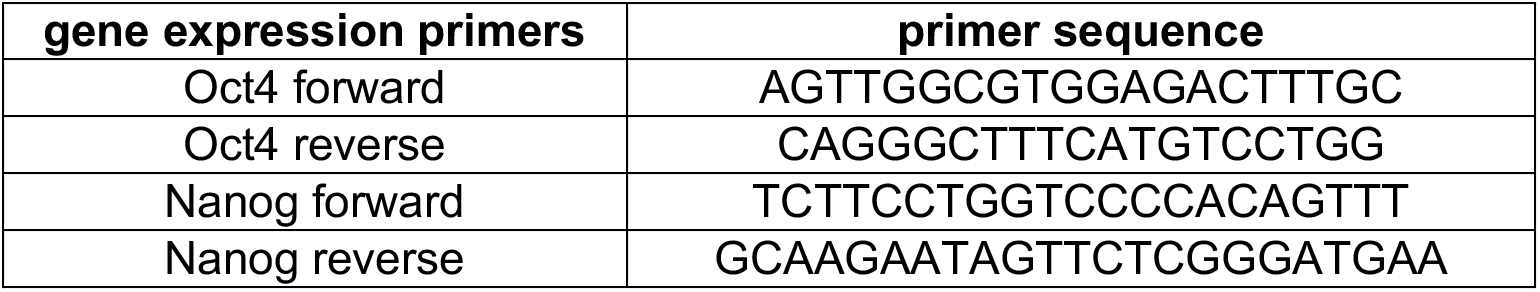

### RNA-seq library preparation and analysis

RNA-seq library preparation was performed using the TruSeq RNA sample preparation kit (Illumina RS-122-2001) as per manufacturer’s instructions. Samples were sequenced on the Illumina HiSeq 2500 platform (Illumina) with 100 bp paired-end reads. RNA-seq reads were aligned to the mouse genome (mm10, NCBI Build 37), RefSeq exons and splicing junctions using BWA alignment algorithm (http://bio-bwa.sourceforge.net/). The reads that were uniquely aligned to the exon and splicing-junction sites for each transcript were then counted as expression level for a corresponding transcript and were subjected to log2 transformation and global median normalization. Differentially expressed genes were identified by the R package DEGseq^40^ using a false discovery rate (FDR) < 0.1, fold change > 1.5.

### Whole Genome BS-Sequencing

The 5mC and 5hmC libraries were prepared using TrueMethyl oxBS Module (NuGEN) following the manufacturer’s protocol. For analysis of the raw data, the MINT pipeline was used^41^. Seq-Monk (https://www.bioinformatics.babraham.ac.uk/projects/seqmonk/) was utilized for visualization.

### Single cell sequencing library generation of E8.5 embryos

*Uhrf1*^+/−^ male and female mice were naturally mated and checked every morning for the appearance of a vaginal plug, the presence of a plug determined the time point E0.5. E8.5 embryos were dissected from the uterus in PBS/FBS and extra-embryonic tissue was removed. The embryos were washed in PBS without Ca^2+^/Mg^2+^ and each was put in a 3 cm dish with 1 ml TrypLE, chopped into small pieces, and incubated at 37°C for 20 min. 300 μl serum was added and pipetted up and down until tissue was dissociated in single cells and filtered. ME-seq libraries were prepared as described in an online protocol (https://www.protocols.io/view/me-seq-protocol-v1-a-single-cell-trimodal-assay-g2wgbyfbx.pdf). Briefly, cells were fixed and permeabilized. For joint measurements of single-cell chromatin accessibility, DNA methylation, and expression, cells were first transposed by Tn5 transposase to mark regions of open chromatin. The mRNA was reverse transcribed using a poly(T) primer containing a unique molecular identifier (UMI) and a random primer. Both primers contained a biotin tag. Permeabilized cells were distributed in a 96-well plate to hybridize well-specific barcoded oligonucleotides to transposed chromatin fragments and poly(T) cDNA. Hybridization was repeated three times to expand the barcoding space and ligate cell barcodes to cDNA and chromatin fragments. Reverse crosslinking was performed to release barcoded molecules. cDNA was separated from chromatin using streptavidin beads, and each library was prepared separately for sequencing. Chromatin DNA was purified, gap-filled and oxidized using mTET1, and deaminated with 1 M pyridine borane before amplification. Both DNA and RNA libraries were quantified with KAPA Library Quantification Kit and pooled to 4 nM for sequencing. Libraries were sequenced on the Illumina NovaSeq platform with 150 PE reads (Read 1: 150 cycles, Index 1: 8 cycles, Index 2: 8 cycles, Read 2: 150 cycles), aiming for a minimum coverage of 50,000 reads per cell.

### Hi-C

Hi-C libraries were prepared using the Arima Hi-C kit from Arima Genomics following the manufacturer’s instructions (A510008). Briefly, ∼400,000 cells from the wild-type or Uhrf1 knockout E8.5 embryos were counted and resuspended in 1 ml of PBS with 3% BSA. Cells were crosslinked using 2% formaldehyde, mixed well by pipetting, and incubated at RT for 10 min. Fixed cells were lysed to release nuclei, and chromatin was digested with proprietary Arima restriction enzymes. DNA overhangs were filled in with biotinylated nucleotides, and spatially proximal fragments were ligated to generate chimeric molecules representative of chromatin contacts. Ligation products were reverse crosslinked, and DNA was purified using AMPure beads. Purified DNA was sonicated on a LE220-plus focused-ultrasonicator (Covaris) for 60 seconds (peak power: 450 W; duty factor: 30%; cycles per burst: 200). Biotinylated proximity-ligation products were size-selected and enriched, then quantified using a Qubit fluorometer. A total of 200 ng of size-selected DNA was used for library preparation with the Arima library preparation kit (A303011), including end repair, A-tailing, adapter ligation, and PCR amplification with indexed primers. Libraries were sequenced on an Illumina NovaSeq platform targeting ∼600 million reads per library. For Hi-C data analysis, FASTQ files were trimmed using Trim Galore! (v2.0) to remove adapters and low-quality bases. Trimmed reads were aligned to the mouse reference genome mm10 using BWA-MEM. Aligned read pairs were parsed, filtered, and deduplicated using pairtools. Contact matrices were generated and balanced using cooler, and downstream visualization and analysis of contact maps were performed using HiContacts. Chromatin loops were called using HiCCUPS with matched parameters across wild-type and Uhrf1 knockout samples.

### CUT&RUN

CUT&RUN was performed using the EpiCypher CUT&RUN kit (14-1048-24rxn) following the manufacturer’s instructions (version 6). A total of 10,000 cells per reaction were used for histone marks H3K4me3 and H3K27me3, and 20,000 cells were used for CTCF. Briefly, ConA beads were activated and bound to cells. Primary antibody was added to each reaction and incubated overnight at 4°C on a nutator. Cells were then incubated with pAG-MNase at room temperature for 10 minutes, followed by targeted chromatin digestion for 2 hours at 4°C. Libraries were prepared using the CUT&RUN library prep kit (14-1001), including end repair, adapter ligation, and PCR amplification with unique i5/i7 indexing primer pairs under the following cycling conditions: 98°C 45 s; 16 cycles of 98°C 15 s, 60°C 10 s, 72°C 60 s; final extension 72°C 60 s; hold at 4°C. Library quality was assessed by FlashGel and Qubit fluorometry. Paired-end sequencing was performed on an Illumina NovaSeq platform. FASTQ files were trimmed of adapters and low-quality sequences using Trim Galore!. Trimmed paired-end reads were aligned to the GRCm38 (mm10) reference genome using Bowtie2 (v2.5.5). BAM files were sorted and indexed with SAMtools, and duplicate reads were marked using Picard (v3.4.0). BedGraph files were generated with bedtools (v2.31.0), and bigWig files were produced using bedGraphToBigWig (v4.8.2) for visualization in the Integrative Genomics Viewer (IGV). Peak calling was performed using MACS2 (v2.2.9.1) for sharp marks and SICER2 (v2.1.0) for broad marks, and peaks were merged into consensus peak sets using bedtools.

### Mass spectrometry-based proteomics

Bulk proteomic profiling was performed using 100,000 cells from wild-type or Uhrf1 knockout E8.5 embryos per assay. Cells were lysed, proteins were extracted and enzymatically digested, and resulting peptides were analyzed by mass spectrometry. Protein abundance matrices were generated from the mass spectrometry data and used for downstream differential analysis. Protein abundance values were normalized and tested for differential abundance between wild-type and Uhrf1 knockout samples using DESeq2, with multiple-testing correction applied to obtain adjusted p values.

### ME-seq sequencing data preprocessing

We developed and applied a versatile pipeline for demultiplexing and mapping sequencing data generated by technologies established in our group. The workflow consists of five main modules: (1) demultiplexing FASTQ files and assigning cell barcode information to each read; (2) adapter trimming and filtering of low-quality reads; (3) read alignment; (4) BAM file processing and quality control; and (5) generation of final gene expression, chromatin accessibility, or methylome profiles. Steps 1 and 2 are identical for sequencing results from multi-omic profiling (scRNA-seq, scATAC-seq, and sc-methyl-seq), whereas steps 3 to 5 are divided into separate branches: ‘a’ transcriptome, ‘b’ methylome, and ‘c’ chromatin accessibility.

Step 1. FASTQ files were generated from base calls using bcl2fastq. Sequencing reads from the Read 2 file were demultiplexed, allowing one mismatch per 8-base barcode across the three rounds of barcoding. Barcode sequences were converted into read identifiers in the format RoundX.xxx (where X denotes the round number and xxx the sequence number) and appended to the header of each read.

Step 2. Reads containing adapter sequence were trimmed and the low-quality reads were removed with Trim Galore using default parameters.

Step 3.a (transcriptome). Reads from step 2 were aligned to the mouse reference genome (mm10) with STAR^42^ using the following settings: --chimOutType WithinBAM --runThreadN 16 --outFilterMultimapNmax 20 --outFilterScoreMinOverLread 0.3 --outFilterMatchNminOverLread 0.3 --outSAMattributes NH HI AS nM MD --limitOutSJcollapsed 5000000 --outSAMtype BAM Unsorted --limitIObufferSize 400000000 --outReadsUnmapped None --readFilesCommand zcat. Step 3.b (methylome and chromatin accessibility). Trimmed reads were aligned to the mouse reference genome (mm10) with Bismark using default settings (Bismark --non_directional-X 2000).

Step 4.a (transcriptome) STAR mapped reads were first filtered with the-F 256 flag to retain uniquely mapped reads and the primary alignment position of multi-mapped reads. To distinguish RNA reads captured by poly(T) primers (dT) from those captured by random RT primers (N6), we used the ‘PT’ tag (PT=1 for dT and PT=0 for N6). The filtered reads were then annotated to both exons and introns using featureCounts^43^. To speed up processing, only barcode combinations with > 100 reads were retained. The filtered and sorted BAM file was convert to h5 file for the next step.

Step 4.b (methylome) Duplicate reads were identified and removed using deduplicate_bismark with the --barcode option. The first and last 9 bases of each read were marked as unmethylated due to the characteristics of Tn5 transposition. Methylated cytosines in CpG (mCG) or non-CpG (mCH) contexts were then identified using bismark_methylation_extractor.

Step 4.c (chromatin accessibility) Duplicates were removed using Picard. Open chromatin region peaks were called on individual samples using MACS2 peak caller^44^ with the following parameters: –nomodel –nolambda –keep-dup-call-summits.

Step 5.a (transcriptome) H5 file from step 4.a was read and processed using Seurat (v5.3.0)^45^. Cells with high ambient RNA contamination (score > 0.5) and doublets were identified and removed using DecontX^46^ and DoubeltFinder^47^. Quality control filtering excluded cells expressing fewer than 800 genes, more than 7,000 genes, or with >3% mitochondrial transcripts. Expression counts were determined by unique read counts to generate a cell-by-gene expression matrix, after which mitochondrial genes were removed. The scRNA-seq data were normalized to counts per 10,000 using Seurat. For visualization, the top 3,000 variable genes were projected into two-dimensional space using UMAP.

Step 5.b (methylome) Tab-delimited (ALLC) files containing methylation levels for every cytosine position was generated from the BED file in step 4. A matrix of peak methylation ratio by cell was created and used as input matrix for Signac^48^.

Step 5.c (chromatin accessibility) Fragment counts within peaks were aggregated to generate a cell-by-peak count matrix. A TF-IDF matrix was then computed from this count matrix using truncated singular value decomposition (RunSVD function, Signac). For visualization, UMAP was performed with the Seurat RunUMAP function, using LSI components 2 through 50.

### Cell-type annotation

We clustered the data and separately annotated the cells with the help of a published study of mouse embryogenesis^49^ using the following label transfer functions of Seurat: FindTransferAnchors and TransferData. Then we annotated the cluster with based on the frequency of transferred cell type annotations.

### Differential expression and differential accessibility analysis

We performed differential gene expression (DEG) and differential accessibility (DA) between wild-type and *Uhrf1* knockout in each cell type. DEGs between groups in each cell type were computed using function FindAllMarkers in Seurat. The Wilcoxon rank-sum test was applied. Genes with fold change higher than 1 and *p* value lower than 0.01 were defined as DEGs. DA between groups of cells was computed by constructing a logistic regression model predicting group membership based on the accessibility of a given peak in the set of cells being compared, with the total number of counts in each cell included as a latent variable in the model and comparing this with a null model using a likelihood ratio test.

### Gene functional enrichment analysis

Gene functional enrichment analysis of GO terms was performed using the enrichGO function from the ClusterProfiler package. To ensure robustness of the results, multiple testing correction was applied using the Bonferroni method, and an adjusted P value threshold of 0.05 was used to define statistical significance.

### Peak-gene *cis*-association

To calculate *cis* peak–gene associations, we considered all ATAC peaks located within a 50 kb window surrounding each annotated transcription start site (TSS). Peak counts or mean CpG methylation within the peaks and gene expression values were used to compute the observed Spearman correlation coefficient (obs) for each peak–gene pair. To estimate background correlations, we applied chromVAR to generate 100 background peaks for each observed peak, matched for accessibility and GC content. Spearman correlations were then calculated between these background peaks and the corresponding gene, producing a null distribution of peak-gene correlation values independent of genomic proximity. For simplicity, a one-sided z-test was used to assign p-values. For peaks associated with multiple genes, only the peak-gene association with the smallest p-value was retained.

### Computing TF footprinting scores

TF binding to DNA protects the occupied binding site from transposition, while the displacement or depletion of adjacent nucleosomes increases accessibility in the surrounding flanking regions. Together, these features constitute the TF footprint. To accurately profile TF footprints, cells of the same type and experimental group (wild-type or Uhrf1 knockout) were aggregated to generate pseudo-bulk ATAC-seq profiles, which were then used for footprinting analysis by ArchR. For each single-base position, we defined a central footprint window flanked by windows on either side and calculated the observed ratio of Tn5 insertion counts as center / (center + flanking). The observed foreground ratio was compared against a background distribution to determine statistical significance, which was subsequently transformed into a footprint score.

### Cell-cell communication

The standard workflow of CellChat (v1.5.0) on single-cell gene expression of ligands and receptors was performed. Based on an existing database of ligand-receptor pairs, we utilized the default parameters ‘mean = trimean’ and ‘trim = 0.1’ to infer a cell-to-cell communication network. Two separate analyses were done for wild-type and knockout conditions.

---

**Extended Data Fig.1.**
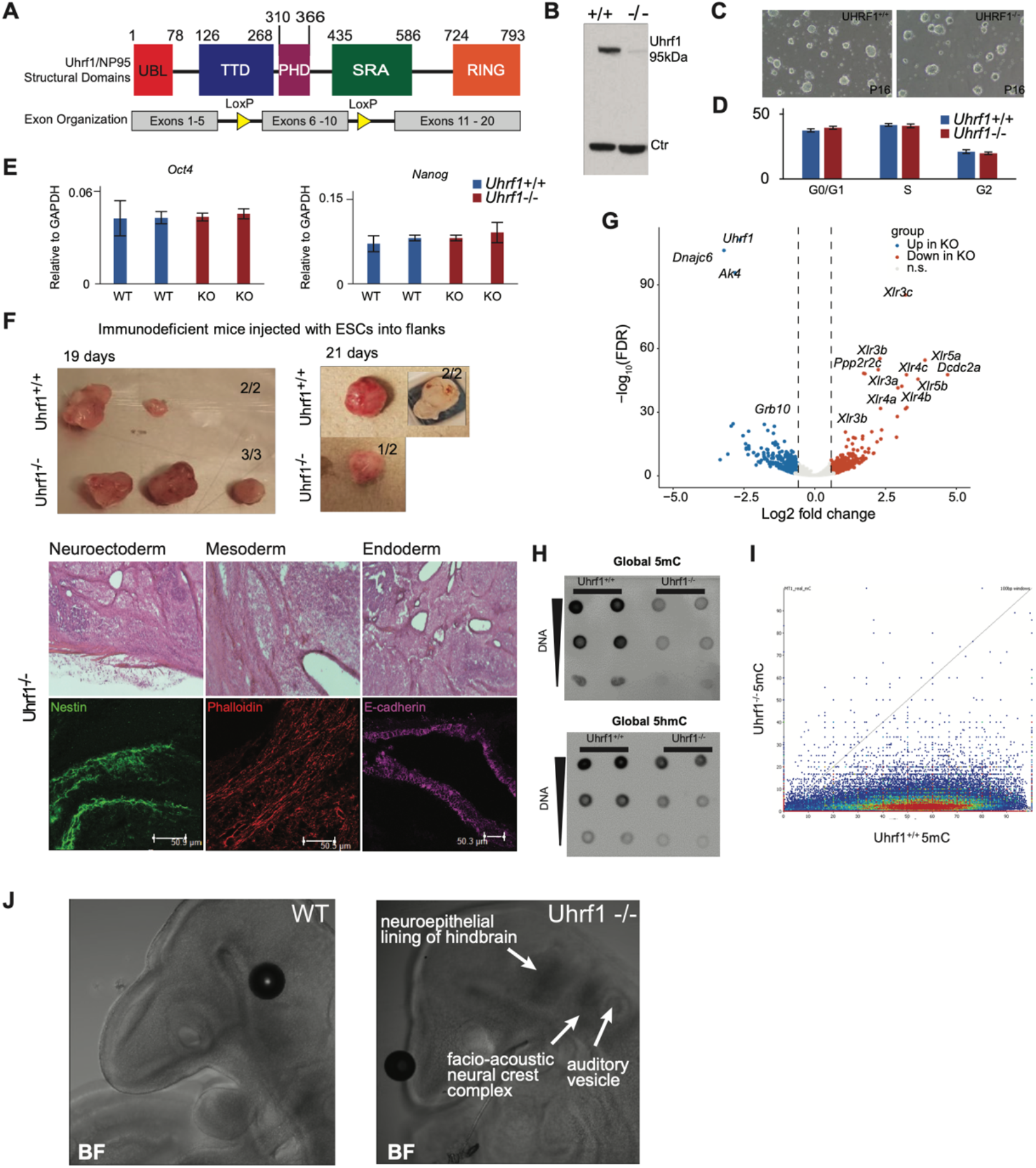
*Uhrf1* knockout does not cause significant phenotypic changes *in vitro*. Related to. Figure 1**. (A)** Schematic of the *Uhrf1* knockout model. **(B)** Western blot confirming reduced UHRF1 protein expression in cultured ESCs. **(C)** Phenotype analysis of *Uhrf1* wild-type and knockout ESCs. **(D)** Cell cycle analysis of *Uhrf1* wild-type and knockout ESCs. **(E)** RT-qPCR of pluripotency markers (Oct4 and Nanog). **(F)** Teratoma assays in immunocompromised mice with lineage-marker staining (Nestin, oectoderm; Phalloidin, mesoderm; E-cadherin, endoderm). **(G)** Volcano plot of differentially expressed genes between wild-type and knockout ESCs. **(H)** Dot blot of 5mC and 5hmC in ESCs. **(I)** Whole-genome bisulfite sequencing of WT and KO ESCs confirming the global reduction of 5mC. **(J)** Brightfield images associated with Figure 1D.

**Extended Data Fig.2.**
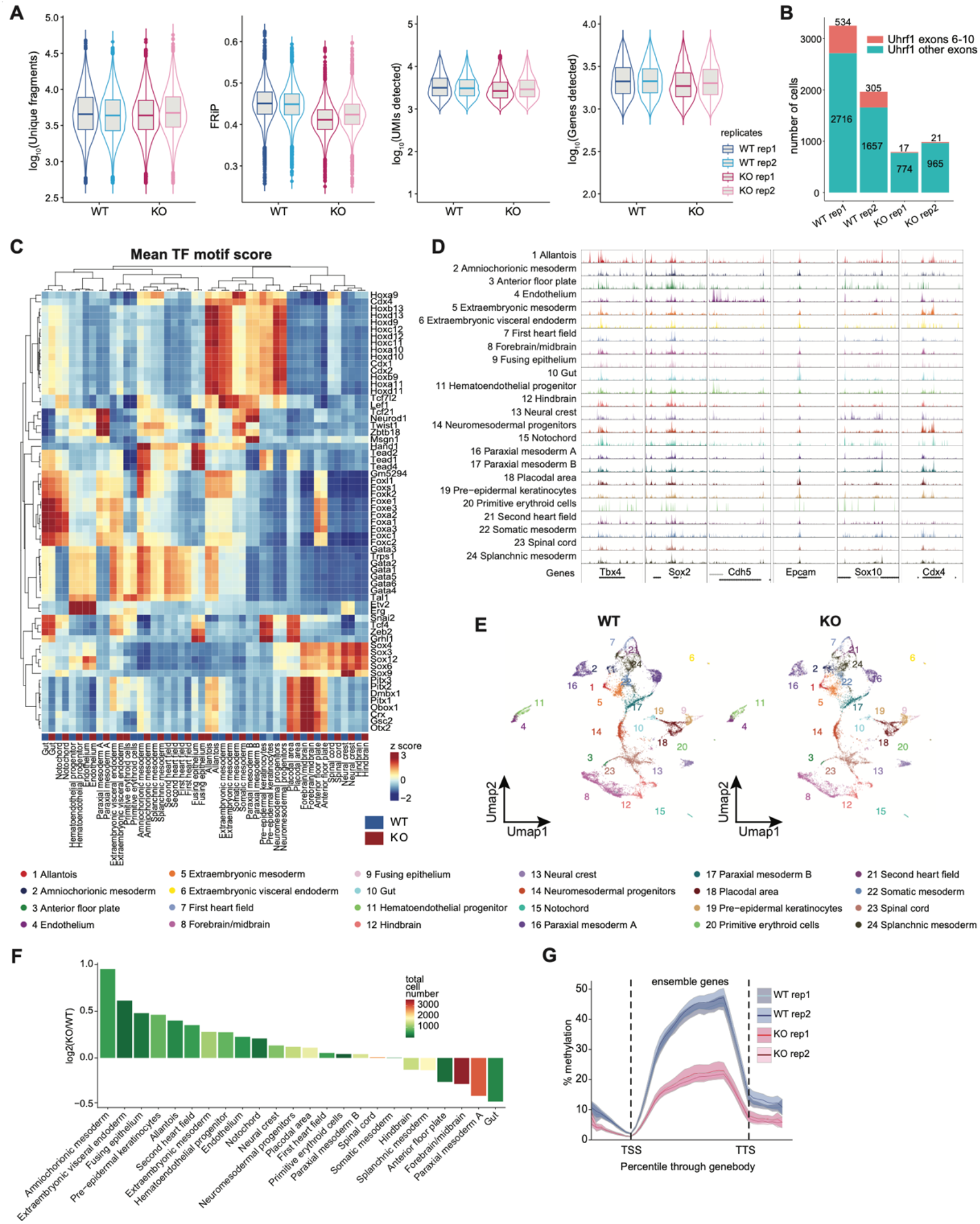
ME-seq generates a high-quality trimodal single-cell atlas capturing cell type composition and methylation changes caused by *Uhrf1* knockout. Related to. Figure 2**. (A)** Quality control of ME-seq data. **(B)** The number of cells detected with *Uhrf1* RNA expression in ME-seq data confirming the deletion of exons 6-10 in knockout embryos. **(C)** Top variable TF motif scores derived from scATAC-seq data in each cell type and condition. **(D)** UMAP colored by the identified cell types, in wild-type and knockout embryos. **(E)** Genomic tracks of scATAC-seq data around representative cell-type-specific marker genes. **(F)** Cell type abundance changes between knockout and wild-type. **(G)** The average DNA methylation across gene bodies was reduced by Uhrf1 knockout.

**Extended Data Fig.3.**
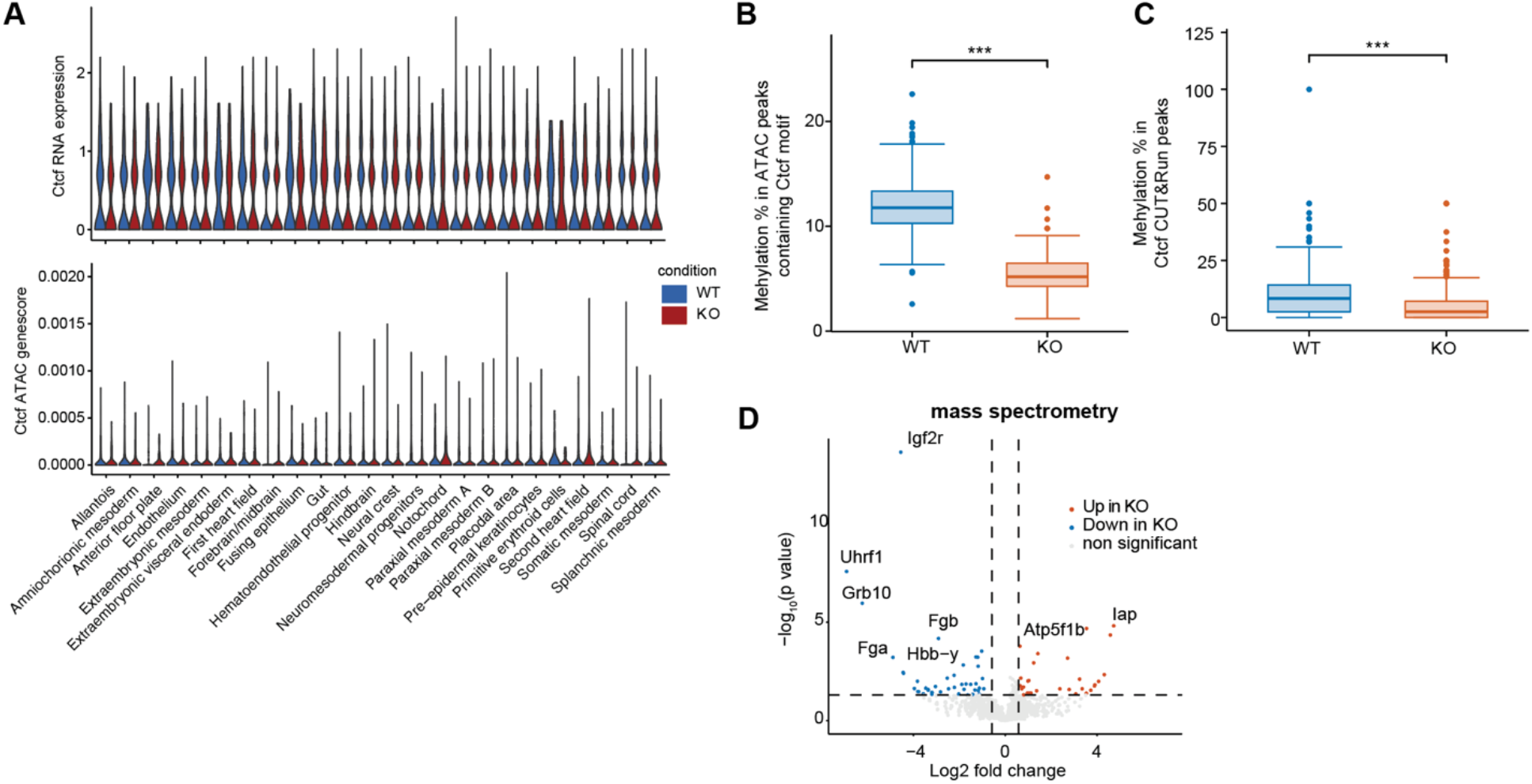
UHRF1 loss reduces DNA methylation at CTCF-associated regulatory regions without altering Ctcf expression or accessibility at the Ctcf locus. Related to. Figure 3**. (A)** Violin plots showing Ctcf RNA expression and chromatin accessibility at the Ctcf locus. **(B)** CpG methylation levels within ATAC peaks containing CTCF motifs in wild-type and UHRF1 knockout embryos. \*\*\**p* < 0.001, Wilcoxon test. **(C)** CpG methylation levels within CTCF CUT&RUN peak regions in wild-type and UHRF1 knockout embryos. \*\*\**p* < 0.001, Wilcoxon test. **(D)** Volcano plot showing protein abundance changes measured by mass spectrometry in wild-type and knockout embryos.

**Extended Data Fig.4.**
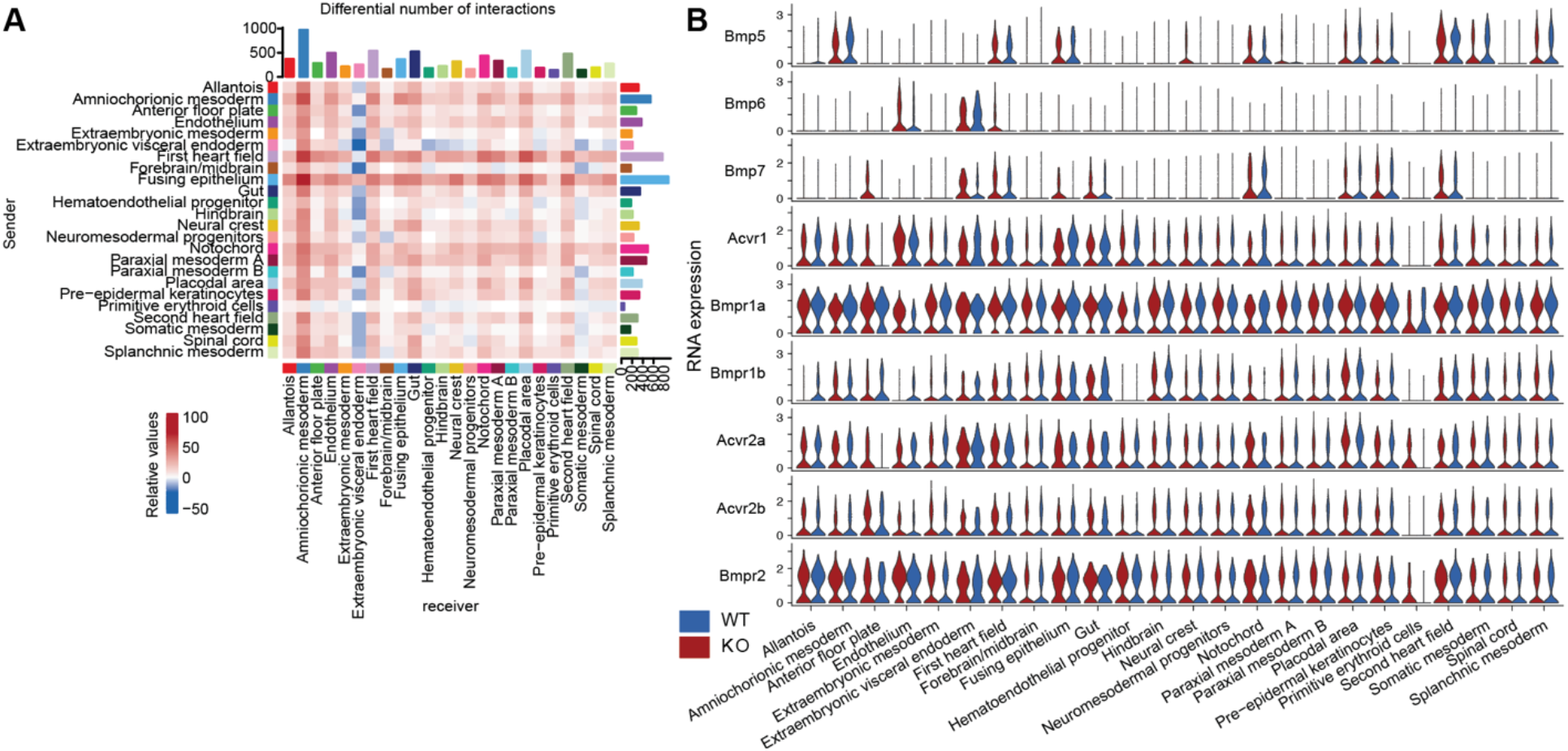
Neural crest cells show reduced predicted communication in UHRF1-null embryos. Related to. Figure 4**. (A)** The differences in the number of cell-cell interactions detected in each cell type. **(B)** Comparing the cell-cell interaction strength between wild-type and knockout across each individual pathway.

**Extended Data Fig.5.**
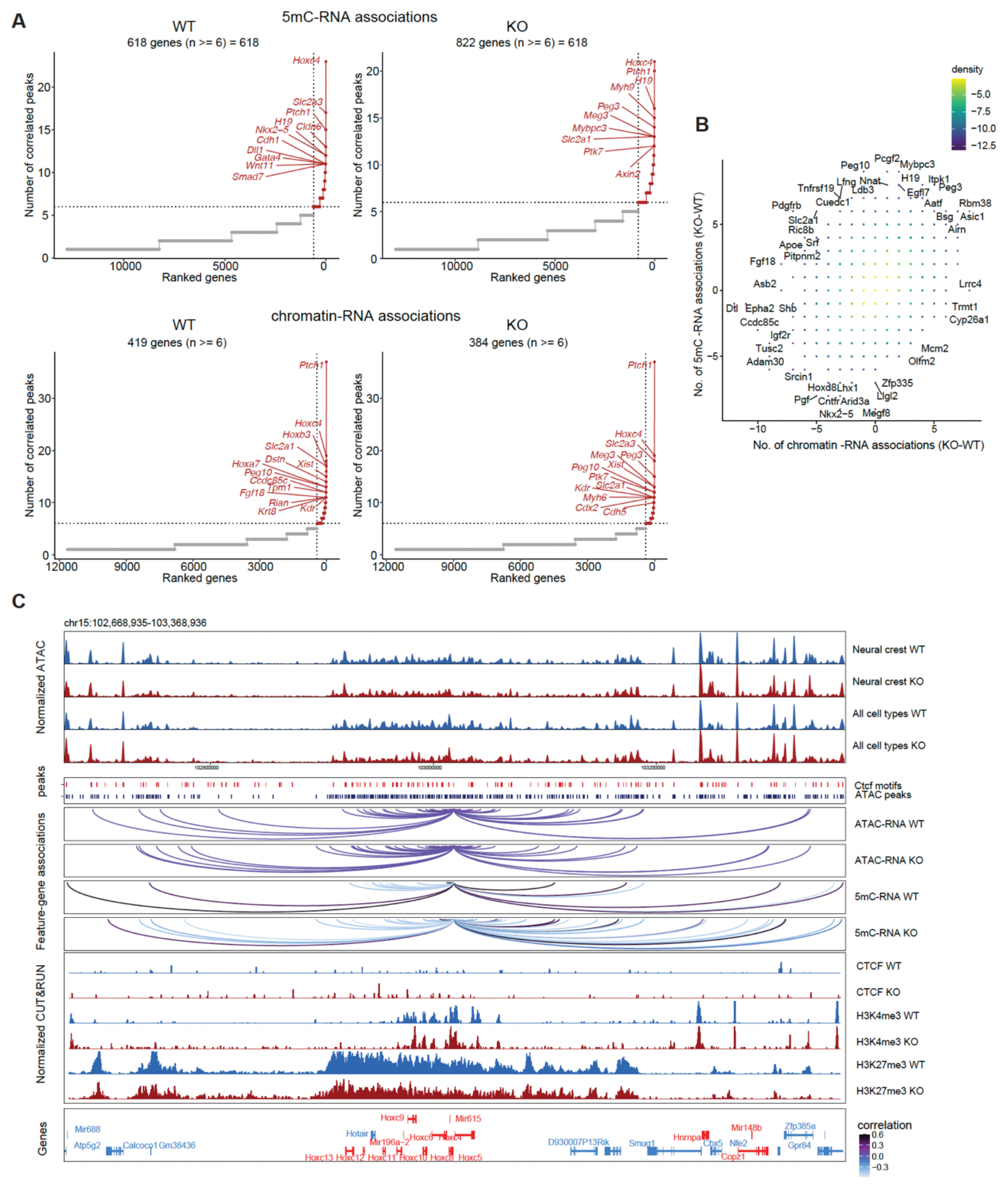
*Uhrf1* loss alters chromatin–RNA relationships associated with DNA methylation. Related to. Figure 5**. (A)** The number of significantly correlated peaks (p < 0.05) for each gene (50 kb from TSSs). The correlation to RNA expression was calculated using DNA methylation or accessibility in peaks. **(B)** The changes in the number of correlated peaks for each gene. **(C)** Genome browser view of the Hoxc loci showing normalized ATAC signal, feature-gene associations, CTCF CUT&RUN, H3K4me3 CUT&RUN, H3K27me3 CUT&RUN, and gene annotations in wild-type and knockout embryos. Loops denote correlations between regulatory features and *Hoxc10* RNA expression, with loop height representing the significance of the correlation and color indicating correlation strength.

**Extended Data Fig. 6.**
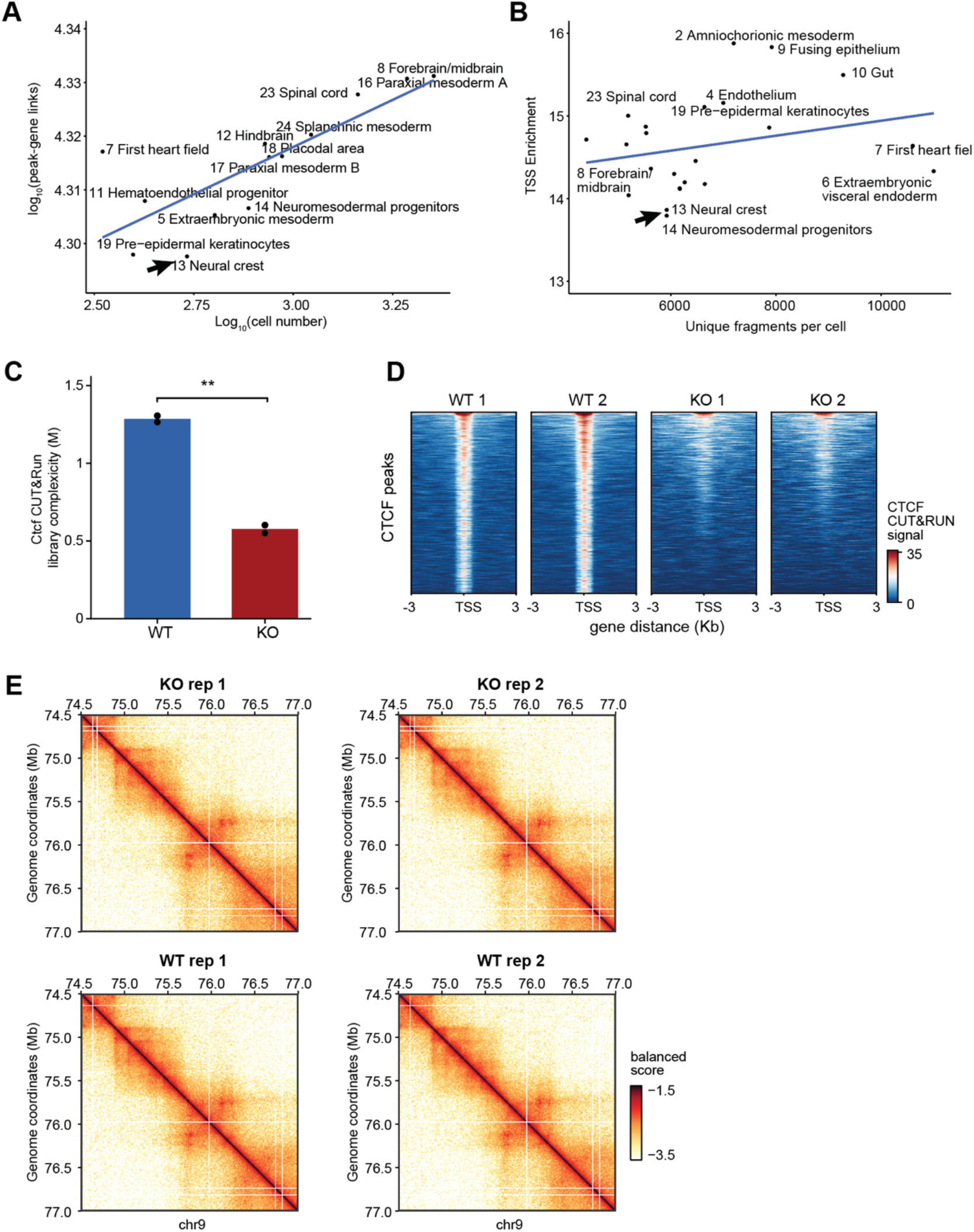
UHRF1 loss is associated with altered chromatin contact organization and reduced CTCF CUT&RUN signal. Related to. Figure 5. **(A)** Relationship between the number of detected peak-gene links and cell number across embryonic cell populations. **(B)** Relationship between TSS enrichment and unique fragments per cell across embryonic cell populations. **(C)** CTCF CUT&RUN library complexity in wild-type and knockout embryos. \*\**p* < 0.01. **(D)** Heatmaps showing CTCF CUT&RUN signal centered at CTCF peaks in biological replicates. **(E)** Representative Hi-C contact maps for wild-type and knockout biological replicates at chr9, showing reproducible chromatin contact patterns across replicates.

## References

1. Unoki, M., Bronner, C. & Mousli, M. A concern regarding the current confusion with the human homolog of mouse Np95, ICBP90/UHRF1. Radiat Res 169, 240–4 (2008).

2. Murao, N. et al. Np95/Uhrf1 regulates tumor suppressor gene expression of neural stem/precursor cells, contributing to neurogenesis in the adult mouse brain. Neurosci Res 143, 31–43 (2019).

3. Bostick, M. et al. UHRF1 plays a role in maintaining DNA methylation in mammalian cells. Science 317, 1760–4 (2007).

4. Rothbart, S.B. et al. Multivalent histone engagement by the linked tandem Tudor and PHD domains of UHRF1 is required for the epigenetic inheritance of DNA methylation. Genes Dev 27, 1288–98 (2013).

5. Vaughan, R.M., Rothbart, S.B. & Dickson, B.M. The finger loop of the SRA domain in the E3 ligase UHRF1 is a regulator of ubiquitin targeting and is required for the maintenance of DNA methylation. J Biol Chem 294, 15724–15732 (2019).

6. Nishiyama, A. et al. Uhrf1-dependent H3K23 ubiquitylation couples maintenance DNA methylation and replication. Nature 502, 249–53 (2013).

7. Wang, L. et al. MOF-mediated acetylation of UHRF1 enhances UHRF1 E3 ligase activity to facilitate DNA methylation maintenance. Cell Rep 43, 113908 (2024).

8. Houliston, R.S. et al. Conformational dynamics of the TTD-PHD histone reader module of the UHRF1 epigenetic regulator reveals multiple histone-binding states, allosteric regulation, and druggability. J Biol Chem 292, 20947–20959 (2017).

9. Choudalakis, M., Kungulovski, G., Mauser, R., Bashtrykov, P. & Jeltsch, A. Refined read-out: The hUHRF1 Tandem-Tudor domain prefers binding to histone H3 tails containing K4me1 in the context of H3K9me2/3. Protein Sci 32, e4760 (2023).

10. Jiang, Q. et al. G9a Plays Distinct Roles in Maintaining DNA Methylation, Retrotransposon Silencing, and Chromatin Looping. Cell Rep 33, 108315 (2020).

11. Kim, A. & Benavente, C.A. Oncogenic Roles of UHRF1 in Cancer. Epigenomes 8(2024).

12. Bell, A.C., West, A.G. & Felsenfeld, G. The protein CTCF is required for the enhancer blocking activity of vertebrate insulators. Cell 98, 387–96 (1999).

13. Wang, H. et al. Widespread plasticity in CTCF occupancy linked to DNA methylation. Genome Res 22, 1680–8 (2012).

14. Roseman, S.A. et al. DNA methylation insulates genic regions from CTCF loops near nuclear speckles. Elife 13(2025).

15. Monteagudo-Sánchez, A., Richard Albert, J., Scarpa, M., Noordermeer, D. & Greenberg, M.V.C. The impact of the embryonic DNA methylation program on CTCF-mediated genome regulation. Nucleic Acids Res 52, 10934–10950 (2024).

16. Yang, Y. et al. Machine learning for classifying tuberculosis drug-resistance from DNA sequencing data. Bioinformatics 34, 1666–1671 (2018).

17. Dahlet, T. et al. Genome-wide analysis in the mouse embryo reveals the importance of DNA methylation for transcription integrity. Nature Communications 11, 3153 (2020).

18. Mulas, C. et al. Defined conditions for propagation and manipulation of mouse embryonic stem cells. Development 146(2019).

19. Sharif, J. et al. The SRA protein Np95 mediates epigenetic inheritance by recruiting Dnmt1 to methylated DNA. Nature 450, 908–912 (2007).

20. Wu, Y. et al. Maternal UHRF1 Is Essential for Transcription Landscapes and Repression of Repetitive Elements During the Maternal-to-Zygotic Transition. Frontiers in Cell and Developmental Biology Volume 8 - 2020(2021).

21. Maenohara, S. et al. Role of UHRF1 in de novo DNA methylation in oocytes and maintenance methylation in preimplantation embryos. PLoS Genet 13, e1007042 (2017).

22. Cao, Y. et al. Deletion of maternal UHRF1 severely reduces mouse oocyte quality and causes developmental defects in preimplantation embryos. The FASEB Journal 33, 8294–8305 (2019).

23. Li, E., Bestor, T.H. & Jaenisch, R. Targeted mutation of the DNA methyltransferase gene results in embryonic lethality. Cell 69, 915–926 (1992).

24. Ma, S. et al. Chromatin Potential Identified by Shared Single-Cell Profiling of RNA and Chromatin. Cell 183, 1103–1116.e20 (2020).

25. Liu, Y. et al. Bisulfite-free direct detection of 5-methylcytosine and 5-hydroxymethylcytosine at base resolution. Nature Biotechnology 37, 424–429 (2019).

26. Pijuan-Sala, B. et al. A single-cell molecular map of mouse gastrulation and early organogenesis. Nature 566, 490–495 (2019).

27. Pachano, T., Haro, E. & Rada-Iglesias, A. Enhancer-gene specificity in development and disease. Development 149(2022).

28. Wang, R.N. et al. Bone Morphogenetic Protein (BMP) signaling in development and human diseases. Genes & Diseases 1, 87–105 (2014).

29. Kartha, V.K. et al. Functional inference of gene regulation using single-cell multi-omics. Cell Genomics 2, 100166 (2022).

30. Qi, C., Di Minin, G., Vercellino, I., Wutz, A. & Korkhov, V.M. Structural basis of sterol recognition by human hedgehog receptor PTCH1. Science Advances 5, eaaw6490 (2019).

31. Zhang, B. et al. Repulsive axon guidance molecule Slit3 is a novel angiogenic factor. Blood 114, 4300–9 (2009).

32. Auvray, C. et al. HOXC4 homeoprotein efficiently expands human hematopoietic stem cells and triggers similar molecular alterations as HOXB4. Haematologica 97, 168–78 (2012).

33. Lister, R. et al. Global epigenomic reconfiguration during mammalian brain development. Science 341, 1237905 (2013).

34. Bronner, C., Alhosin, M., Hamiche, A. & Mousli, M. Coordinated Dialogue between UHRF1 and DNMT1 to Ensure Faithful Inheritance of Methylated DNA Patterns. Genes (Basel) 10(2019).

35. Schmidt, C.S. et al. Global DNA Hypomethylation Prevents Consolidation of Differentiation Programs and Allows Reversion to the Embryonic Stem Cell State. PLOS ONE 7, e52629 (2012).

36. Imaz-Rosshandler, I. et al. Tracking early mammalian organogenesis – prediction and validation of differentiation trajectories at whole organism scale. Development 151(2024).

37. Kunath, T. et al. FGF stimulation of the Erk1/2 signalling cascade triggers transition of pluripotent embryonic stem cells from self-renewal to lineage commitment. Development 134, 2895–2902 (2007).

38. Betschinger, J. et al. Exit from pluripotency is gated by intracellular redistribution of the bHLH transcription factor Tfe3. Cell 153, 335–47 (2013).

39. Nichols, J., Silva, J., Roode, M. & Smith, A. Suppression of Erk signalling promotes ground state pluripotency in the mouse embryo. Development 136, 3215–3222 (2009).

40. Wang, L., Feng, Z., Wang, X., Wang, X. & Zhang, X. DEGseq: an R package for identifying differentially expressed genes from RNA-seq data. Bioinformatics 26, 136–8 (2010).

41. Cavalcante, R.G., Patil, S., Park, Y., Rozek, L.S. & Sartor, M.A. Integrating DNA Methylation and Hydroxymethylation Data with the Mint Pipeline. Cancer Res 77, e27–e30 (2017).

42. Dobin, A. et al. STAR: ultrafast universal RNA-seq aligner. Bioinformatics 29, 15–21 (2013).

43. Liao, Y., Smyth, G.K. & Shi, W. featureCounts: an efficient general purpose program for assigning sequence reads to genomic features. Bioinformatics 30, 923–30 (2014).

44. Zhang, Y. et al. Model-based analysis of ChIP-Seq (MACS). Genome Biol 9, R137 (2008).

45. Stuart, T. et al. Comprehensive Integration of Single-Cell Data. Cell 177, 1888–1902.e21 (2019).

46. Yang, S. et al. Decontamination of ambient RNA in single-cell RNA-seq with DecontX. Genome Biol 21, 57 (2020).

47. McGinnis, C.S., Murrow, L.M. & Gartner, Z.J. DoubletFinder: Doublet Detection in Single-Cell RNA Sequencing Data Using Artificial Nearest Neighbors. Cell Syst 8, 329–337.e4 (2019).

48. Stuart, T., Srivastava, A., Madad, S., Lareau, C.A. & Satija, R. Single-cell chromatin state analysis with Signac. Nat Methods 18, 1333–1341 (2021).

49. Qiu, C. et al. Systematic reconstruction of cellular trajectories across mouse embryogenesis. Nat Genet 54, 328–341 (2022).

